# Mapping the substrate landscape of protein phosphatase 2A catalytic subunit PPP2CA

**DOI:** 10.1101/2023.09.19.558429

**Authors:** Abigail Brewer, Gajanan Sathe, Billie E. Pflug, Thomas J. Macartney, Gopal P. Sapkota

## Abstract

Protein phosphatase 2A (PP2A) is an essential Ser/Thr phosphatase that regulates a plethora of cellular processes. PP2A operates as a holoenzyme complex, comprising one each of the scaffolding (A), regulatory (B) and catalytic (C) subunits. PPP2CA is the principal catalytic subunit of the PP2A holoenzyme complex. Although previous studies have reported many substrates of specific PP2A holoenzyme complexes, the full scope of PP2A substrates in cells remains to be defined. To address this, we generated HEK293 cells in which PPP2CA was homozygously knocked in with a dTAG, allowing for efficient and selective degradation of dTAG-PPP2CA with proteolysis-targeting chimeras (PROTACs) targeting the dTAG. By employing an unbiased global phospho-proteomic analysis, we identified 6,280 phospho-peptides corresponding to 2,204 proteins that showed a significant increase in abundance upon dTAG-PPP2CA degradation, implicating them as potential PPP2CA substrates. Among these, some were established PP2A substrates, while most were novel. Bioinformatic analyses revealed the involvement of the identified potential PPP2CA substrates in many cellular processes, including spliceosome function, the cell cycle, RNA transport and ubiquitin-mediated proteolysis. We show that a pSP/pTP motif is a predominant target for PPP2CA. We confirmed some of our phospho-proteomic data with immunoblotting, by utilising commercially available phospho-specific antibodies. We provide an in-depth atlas of potential PPP2CA substrates and establish targeted degradation as a robust tool to unveil phosphatase substrates in cells.

## Introduction

Protein phosphorylation is a fundamental post-translational modification that controls virtually every cellular process. Phosphorylation is a reversible event comprising the addition of a phosphate group through formation of a hydrolysable phospho-ester bond. In proteins, this modification occurs primarily on serine, threonine, or tyrosine residues. In response to specific signals, phosphorylation can alter protein conformation, stability, catalytic activity, subcellular localisation, or interactions with other partners, thereby altering protein function, cell signalling and, ultimately, cellular fate decisions (Cohen, 2001). Protein kinases catalyse the attachment of phosphate to target proteins while phosphatases elicit hydrolysis to remove the phosphate. Balancing the activities of these enzymes enables fine-tuning of protein phosphorylation states and consequently these enzymes serve crucial roles in metabolism, maintaining homeostasis, cellular transport and secretory processes. Unsurprisingly, numerous human diseases are associated with aberrant regulation of protein phosphorylation, including many cancers and neurodegenerative disorders (Perluigi et al., 2016, Singh et al., 2017, Sontag, 2001). For example, hyper-activation of human epidermal growth factor receptor 2 (HER2) kinase, which leads to increased mitogen-activated protein kinase (MAPK) pathway signalling and proliferation, has been linked to breast cancer (Neve et al., 2002). Similarly, hyperphosphorylation of tau protein by kinases such as cyclin-dependent kinase 5 (CDK5) and glycogen synthase kinase 3β (GSK3β) has been implicated in the formation of neurofibrillary tangles that have been associated with development of Alzheimer’s disease (Plattner et al., 2006). As such, protein kinases and phosphatases have been explored as potential drug targets. While many specific kinase inhibitors have been developed, with around 75 approved for clinical use by the US Food and Drug Administration (FDA) (Druker et al., 2001, Cohen et al., 2021, Lui et al., 2022), equivalent progress has not been seen in the targeting of phosphatases. This is perceived to be a result of the lack of substrate specificity displayed by phosphatases (Kohn, 2020). Nonetheless, the full substrate landscape for different phosphatases remains poorly defined. More than two thirds of the 518 kinases in the human kinome conduct phosphorylation of serine and threonine residues, attributing to around 98% of documented phosphorylation events and forming one of the most common cellular post-translational modifications (Olsen et al., 2006). Yet the majority of serine and threonine dephosphorylation is performed by only two phosphatases: protein phosphatase 1 (PP1) and protein phosphatase 2A (PP2A) (Shi, 2009).

PP2A is an essential Ser/Thr phosphatase that belongs to the PPP (phosphoprotein phosphatase) family. PP2A has been implicated in a broad range of cellular processes such as proliferation, DNA repair, RNA splicing and apoptosis, and is widely considered to display tumour suppressor functions (Eichhorn et al., 2009). Indeed, activation of PP2A through PP2A-activating drugs (PADs) is being investigated as a potential novel drug mechanism to tackle some cancers as well as neurodegenerative and inflammation-mediated diseases (Clark and Ohlmeyer, 2019). Although PP2A is reported to exist in a dimeric complex, comprising a scaffolding A subunit and catalytic C subunit, there is overwhelming evidence that it exists as a trimeric holoenzyme complex where a regulatory B subunit is also involved (Figure S1) (Bryant et al., 1999, Xing et al., 2006, Tolstykh et al., 2000, Cho and Xu, 2007). The trimeric holoenzyme configuration functions to regulate PP2A phosphatase activity with the B subunit providing temporal and spatial selectivity (Zolnierowicz et al., 1994, Sandal et al., 2021). The regulatory B subunits are encoded by 15 different genes and have at least 26 different transcript and splice variants, which are classified into four families: B/B55, B’/B56, B’’/PR72 and B’’’/Striatin (Eichhorn et al., 2009). The B subunits provide substrate selectivity for PP2A through assembly of over 70 possible holoenzyme complexes (Janssens et al., 2008). In contrast to the B subunits, there are only two isoforms each of the scaffolding A subunit (PR65A and B) and the catalytic C subunit (PPP2CA and B). The PPP2CA and PPP2CB isoforms of the catalytic subunit share 97% homology, with variation occurring at the N-terminus (Khew-Goodall et al., 1991). Nonetheless, PPP2CA is typically over ten-fold more abundant than PPP2CB in most cells due to stronger promoter activity and differences in messenger RNA (mRNA) turnover rates (Khew-Goodall and Hemmings, 1988). Interestingly, null mutation of PPP2CA results in early embryonic lethality, indicating that, despite high sequence similarity, PPP2CB is unable to compensate for the loss of PPP2CA (Gu et al., 2012). This suggests that non-redundant functions exist for PPP2CA and PPP2CB, at least in the context of embryonic development.

In addition to being regulated by the formation of the holoenzyme complex, PPP2CA catalytic activity is also tightly controlled through post-translational modification of the C-terminus by phosphorylation and methylation, which serve to promote or impede interaction with regulatory B subunits (Janssens et al., 2008). Furthermore, the presence of similar short sequences, known as short-linear interaction motifs (SLiMs), within PP2A substrates has been identified to promote interaction with specific regulatory B subunits for their dephosphorylation by the holoenzyme complex. The SLiM motif LxxIxE, particularly LSPIxE, has been implicated in binding B56α subunits (Hertz et al., 2016, Wang et al., 2016, Kruse et al., 2020), while p[ST]-P-x(4,10)-[RK]-V-x-x-[VI]-R, found in PP2A substrates such as tau, was associated with B55α engagement (Fowle et al., 2021).

To explore the extent of PP2A holoenzyme activity and define properties of putative PP2A substrates, some phospho-proteomic studies have been reported. One such study identified phospho-proteins from calyculin A-treated HeLa cell extracts that were lost upon incubation of the extracts with recombinant PPP2CA *in vitro* (Hoermann et al., 2020). From this study, it was suggested that PPP2CA intrinsically favours dephosphorylation of phospho-Thr residues. Another study employed an *in vitro* assay system called MRBLE:Dephos to ascertain PP2A-B55 and PP1 amino acid preferences before uncovering putative PP2A-B55 substrates in mitotic exit using phospho-proteomics (Hein et al., 2023). Phospho-proteomic analysis upon short interfering RNA (siRNA)-mediated depletion of PP2A inhibitors, such as cancerous inhibitor of protein phosphatase 2A (CIP2A), SET nuclear proto-oncogene (SET) and protein phosphatase methylesterase 1 (PME-1), or the A scaffolding subunit PPP2R1A, resulted in identification of putative PP2A substrates predicted to be involved in many cellular processes, including RNA splicing, kinase signalling and DNA repair (Kauko et al., 2020). Other studies have explored PP2A substrates controlled by specific holoenzyme configurations or B subunit isoforms (such as PPP2R2A/B55α (Panicker et al., 2020), PP2A-B55/B, - B56/B’ and -PR48/B’’ (Jong et al., 2020), PP2A-B55/B and PP2A-B56/B’ (Kruse et al., 2020), PPP2CB (Gao et al., 2022) and PP2A-Cdc55 in *Saccharomyces cerevisiae* (Baro et al., 2018)), or by probing PP2A activity in response to specific stimuli, such as downstream of modulation of PP2A inhibitors/activators (Bernal et al., 2014). An approach involving the disruption of the core PPP2CA catalytic subunit *in cellulo*, followed by an unbiased phospho-proteomic analysis, could enable delineation of the full range of PPP2CA substrates within that cellular context. However, due to the essential function of PPP2CA for cell survival, its sustained depletion, for example with clustered regularly interspaced short palindromic repeats (CRISPR)/Cas9 genome editing, is not feasible.

Inducible protein degradation can overcome the limitations of prolonged disruption of essential genes and has been employed to interrogate phosphatase function, for example through auxin-mediated degradation of the catalytic subunit of Ser/Thr phosphatase PP6 (PP6c) (Mariano et al., 2023). Recent advances in the targeted protein degradation field have enabled the efficient and selective acute degradation of proteins of interest (POIs) through small heterobifunctional molecules, known as proteolysis-targeting chimeras (PROTACs) (Bondeson et al., 2015, Bondeson and Crews, 2017). PROTACs harness endogenous cellular degradation machinery, such as the ubiquitin proteasome system, thus avoiding the exogenous TIR1 (transport inhibitor response 1) expression, as is required for the auxin-inducible degradation system. By tagging POIs endogenously with degron tags, such as the dTAG (FKBP12^F36V^), using CRISPR/Cas9 genome editing, PROTACs directed at the dTAG, such as dTAG-13 (Nabet et al., 2018), dTAG^V^-1 (Nabet et al., 2020) and dTAG-VHL (Simpson et al., 2022), can lead to the degradation of POIs. Here, we employ dTAG-PROTAC technology to target the degradation of dTAG-PPP2CA, which was knocked-in homozygously to human embryonic kidney (HEK) 293 cells using CRISPR/Cas9 genome editing, before employing unbiased phospho-proteomic analysis to reveal the full repertoire of putative PPP2CA substrates in this cellular context.

## Results

### Generation of ^dTAG/dTAG^PPP2CA knock-in HEK293 cells and assessment of PROTAC-mediated degradation

To explore the substrates of PPP2CA, we generated ^dTAG/dTAG^PPP2CA knock-in HEK293 cells in which dTAG was homozygously inserted at the N-terminus of PPP2CA by using CRISPR/Cas9 technology (Ran et al., 2013) (Fig. S2A-F). Successful knock-in of dTAG was confirmed by a combination of immunoblotting with anti-PPP2CA/B and anti-dTAG antibodies (Fig. 1A and S2C) as well as polymerase chain reaction (PCR) amplification and genomic sequencing of the target gene locus (Fig. S2D-F). A cross-reacting band at ∼48 kDa, the expected molecular weight for dTAG-PPP2CA, was detected by both anti-dTAG and anti-PPP2CA/B immunoblotting of extracts from the ^dTAG/dTAG^PPP2CA knock-in HEK293 cell clone, which was absent in wild type (WT) HEK293 cells, confirming the knock-in (Fig. 1A and S2C). The band at ∼35 kDa observed with anti-PPP2CA/B antibody represents the endogenous PPP2CA and PPP2CB, since the two isoforms share 100% sequence homology for the epitope that the antibody was raised against (residues 289 – 307 of human PPP2CA). Consistent with this, a reduction in the intensity of this 35 kDa band was observed with anti-PPP2CA immunoblotting in ^dTAG/dTAG^PPP2CA HEK293 cells compared to WT HEK293 cells (Fig. 1A and S2C). To ensure homozygous knock-in, the genomic locus around ^dTAG/dTAG^PPP2CA from clone 1 knock-in cells was amplified by PCR and yielded one product at the expected ∼2800 bp molecular weight, compared to ∼1200 bp as was observed in WT cells (Fig. S2D). The PCR product from ^dTAG/dTAG^PPP2CA cells was sequenced and confirmed homozygous knock-in at the expected locus (Fig. S2E-F).

**Figure 1.**
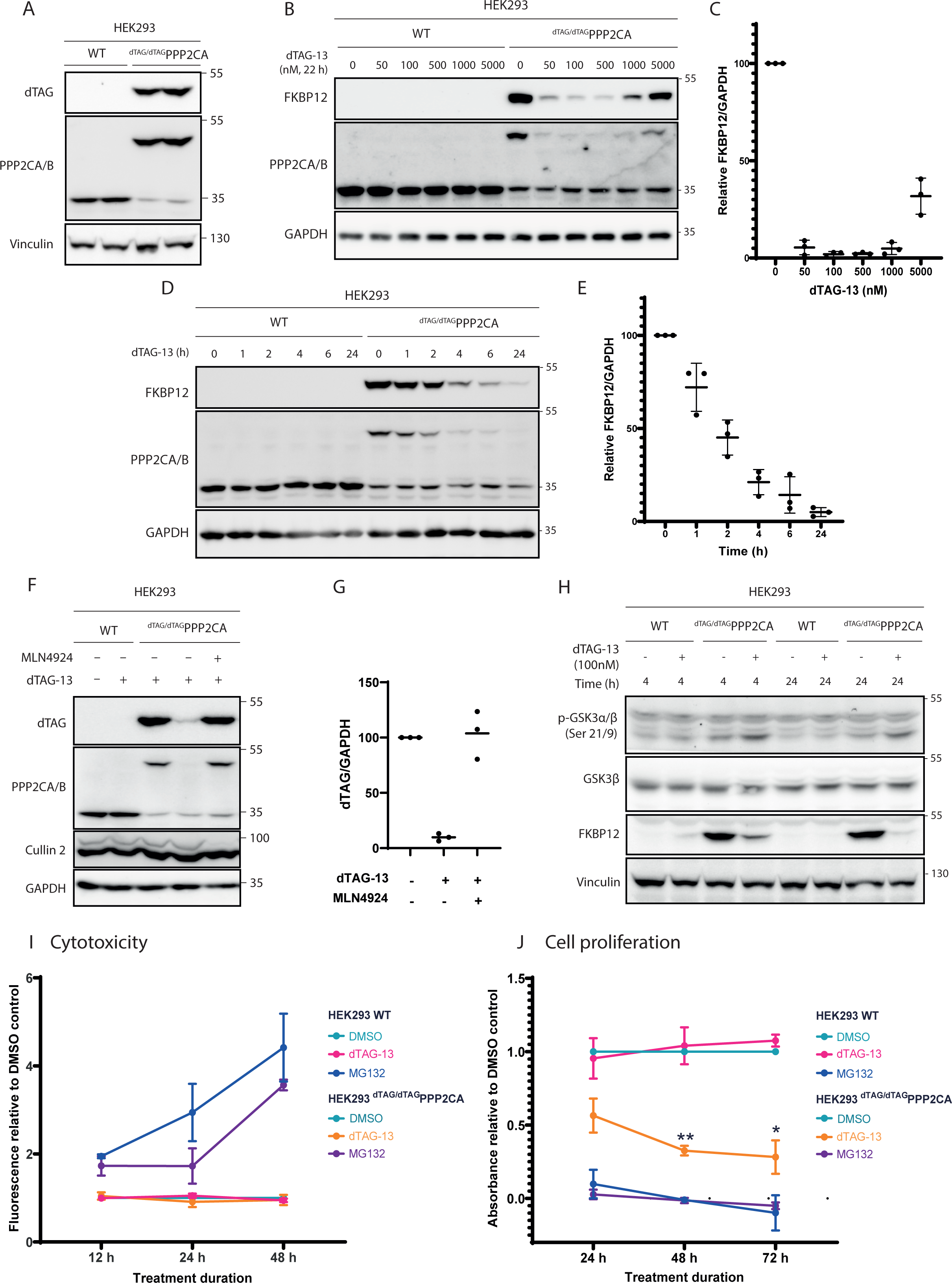
Degradation of dTAG-PPP2CA with dTAG-13. **A.** Characterisation of ^dTAG/dTAG^PPP2CA HEK293 knock-in cell lines. Extracts (20 μg protein) from wild type (WT) and ^dTAG/dTAG^PPP2CA HEK293 cells were resolved by SDS-PAGE and transferred to nitrocellulose membranes, which were analysed by immunoblotting with the indicated antibodies. **B-C.** Dose response of dTAG-13-mediated dTAG-PPP2CA degradation. As in A, except extracts (20 μg protein) were from WT and ^dTAG/dTAG^PPP2CA HEK293 cells treated for 22 h with the indicated concentrations of dTAG-13 or DMSO. Relative dTAG-PPP2CA levels from immunoblots (B) were quantified and are presented in (C) as mean values ± SD from n=3 independent experiments. Anti-FKBP12 antibody was used to detect dTAG (FKBP12^F36V^). **D-E.** Time course of dTAG-13-mediated dTAG-PPP2CA degradation. As in B-C, except extracts (20μg protein) were from WT and ^dTAG/dTAG^PPP2CA HEK293 cells treated with 100 nM dTAG-13 for the indicated time durations. **F-G.** Degradation of dTAG-PPP2CA in ^dTAG/dTAG^PPP2CA HEK293 cells using optimised conditions. As in B-C, except extracts (20μg protein) were from ^dTAG/dTAG^PPP2CA HEK293 cells treated with DMSO, dTAG-13 (100 nM) or a combination of dTAG-13 (100 nM) and MLN4924 (1000 nM) for 24 h prior to lysis. **H.** Confirmation that dTAG-PPP2CA retains phosphatase activity following dTAG knock-in. As in A, except extracts (20μg protein) were from WT and ^dTAG/dTAG^PPP2CA HEK293 cells treated with DMSO or dTAG-13 (100 nM) for 4 h or 24 h prior to lysis. **I.** Cell cytotoxicity assay to determine cytotoxicity of dTAG-13-mediated dTAG-PPP2CA degradation in ^dTAG/dTAG^PPP2CA and WT HEK293 cells. Cells were treated for 12, 24 or 48 h with dTAG-13 (100 nM), DMSO as a negative control or MG132 (40 µM) as a positive control. Cytotoxicity was assessed by using CellTox Green Assay (Promega), with fluorescence being measuring using a PHERAstar plate reader (ex: 480 nm em: 530 nm). Data represent n = 3. Values are shown as a mean fluorescence reading normalized to DMSO controls ± SD. **J.** Cell proliferation assay to determine impact of dTAG-13-mediated dTAG-PPP2CA degradation on cell proliferation. Proliferation was measured using CellTiter 96® AQ_ueous_ One Solution Cell Proliferation Assay (Promega) by treating ^dTAG/dTAG^PPP2CA and WT HEK293 cells with dTAG-13 (100 nM) for 24, 48 or 72 h, with DMSO as a negative control or MG132 (40 μM) as a positive control. Cells were then incubated with the CellTiter 96® AQ_ueous_ One Solution Reagent before absorbance was measured at 490 nm. Data represent n = 3, with 3 technical replicates included per condition for each separate biological repeat. Values are shown as a mean absorbance reading normalized to DMSO controls ± SD, with 2-way ANOVA and Tukey’s HSD post-hoc test used for statistical analysis.

Next, we sought to explore the targeted degradation of dTAG-PPP2CA with dTAG-13 (Nabet et al., 2018). WT and ^dTAG/dTAG^PPP2CA HEK293 cells were treated with increasing concentrations of dTAG-13 (from 50-5000 nM) or dimethylsulphoxide (DMSO) for 22 h. In comparison to DMSO controls, a dose-dependent degradation of dTAG-PPP2CA was observed from 50-500 nM dTAG-13 in ^dTAG/dTAG^PPP2CA HEK293 cells with almost complete degradation observed with dTAG-13 treatment at 100 and 500 nM (Fig. 1B and 1C). At 1000 and 5000 nM dTAG-13, the degradation of dTAG-PPP2CA was less efficient (Fig. 1B), implying a hook effect, which is reminiscent of PROTACs at high treatment concentrations (Pettersson and Crews, 2019). Importantly, dTAG-13 treatment, even at the highest concentration of 5000 nM, did not elicit any change in the abundance of endogenous PPP2CA in WT HEK293 cells (Fig. 1B). A time course experiment in which WT and ^dTAG/dTAG^PPP2CA HEK293 cells were treated with 100 nM dTAG-13 showed a time-dependent degradation of dTAG-PPP2CA, with ∼50% degradation relative to DMSO controls observed at 2 h and almost complete degradation observed at 24 h (Fig. 1D and 1E). Again, no changes in the abundance of endogenous PPP2CA were observed in WT HEK293 cells at any time point following dTAG-13 treatment (Fig. 1D). In comparison to other dTAG-targeting PROTACs, including dTAG-^V^1 and dTAG-VHL, dTAG-13 achieved optimal degradation of dTAG-PPP2CA with the lowest PROTAC concentration, so we continued with dTAG-13 (Figure S3A-D). dTAG-13 recruits dTAG to the cullin 4A (CUL4A)-ring-box 1 (RBX1) E3 ligase complex via the CUL4A substrate receptor cereblon (CRBN) for degradation by the proteasome (Nabet et al., 2018). Consistent with this, treatment of ^dTAG/dTAG^PPP2CA HEK293 cells with the NEDD8 activating enzyme E1 subunit 1 (NAE1) inhibitor MLN4924, which prevents the NEDDylation and activation of all cullins (Soucy et al., 2009), resulted in the rescue of dTAG-PPP2CA degradation caused by dTAG-13 (Fig. 1F and 1G). Compared to DMSO treatment, MLN4294 treatment led to a collapse of the upper cullin 2 (CUL2) band corresponding to NEDDylated species, confirming inhibition of NAE1 (Fig. 1F and 1G).

We also set out to test whether the dTAG knock-in of PPP2CA had any impact on its phosphatase catalytic activity against a reported PP2A substrate, namely p-Ser9 GSK3β (Cross et al., 1995, Shaw et al., 1997, Mitra et al., 2012, Wang et al., 2015). In DMSO-treated WT and ^dTAG/dTAG^PPP2CA HEK293 cells over 4 and 24 h duration, no substantial change in the abundance of p-Ser9 GSK3β was observed between the cell lines, suggesting that the dTAG knock-in on PPP2CA had no effect on the basal dephosphorylation of p-Ser9 GSK3β. The degradation of dTAG-PPP2CA with dTAG-13 in ^dTAG/dTAG^PPP2CA HEK293 cells would be predicted to enhance the abundance of p-Ser9 GSK3β. Indeed, dTAG-13 treatment of ^dTAG/dTAG^PPP2CA HEK293 cells for both 4 h and 24 h caused a marked degradation of dTAG-PPP2CA and concurrently led to enhanced levels of p-Ser9 GSK3β compared to DMSO-treated controls (Fig. 1H). In WT HEK293 cells, no differences in p-Ser9 GSK3β levels were apparent with either DMSO or dTAG-13 treatment at both durations (Fig. 1H). GSK3β protein abundance was unaffected by the dTAG-PPP2CA knock-in or the treatments with either DMSO or dTAG-13.

We then explored whether targeted degradation of dTAG-PPP2CA impacted cell viability and proliferation. We observed no significant cell toxicity up to 48 h following dTAG-13 treatment, in either WT or ^dTAG/dTAG^PPP2CA HEK293 cells (Fig. 1I). However, and perhaps unsurprising, prolonged degradation of dTAG-PPP2CA with dTAG-13 treatment for 48 h and 72 h significantly impeded cell proliferation (Fig. 1J), consistent with previous reports of the importance of PPP2CA to cell function (Gu et al., 2012, Pan et al., 2015, Eichhorn et al., 2009, Wlodarchak and Xing, 2016). Collectively, these data reveal optimal conditions for the dTAG-13-mediated degradation of dTAG-PPP2CA in ^dTAG/dTAG^PPP2CA HEK293 cells and support that this inducible, targeted degradation could enable dissection of the plethora of substrates whose phospho-regulation is controlled by PPP2CA.

### A global total- and phospho-proteomic analysis to elucidate PPP2CA targets

To uncover PPP2CA targets in ^dTAG/dTAG^PPP2CA HEK293 cells, an unbiased tandem mass tag (TMT)-based quantitative proteomic and phospho-proteomic workflow was adopted following degradation of dTAG-PPP2CA with 100 nM dTAG-13 treatment for 24 h (Figure S4). DMSO-treated cells were included as a negative control. Immunoblot analysis of extracts from 3 biological replicates, which were prepared for proteomic and phospho-proteomic analysis, demonstrated robust degradation of dTAG-PPP2CA in ^dTAG/dTAG^PPP2CA HEK293 cells upon treatment with dTAG-13 for 24 h in comparison to DMSO-treated controls (Fig. 2A). Total quantitative proteomic analysis of DMSO- and dTAG-13-treated ^dTAG/dTAG^PPP2CA HEK293 cell extracts identified a total of 80,998 peptides, belonging to 7,589 unique proteins (full list available in Table S1). Of these, the only protein whose abundance was significantly reduced (and with fold change <0.5) in dTAG-13-treated cells compared to DMSO-treated controls was PPP2CA (Fig. 2B), demonstrating the remarkable selectivity of targeted degradation caused by dTAG-13. The quantitative total proteomic data demonstrating that PPP2CA was the only protein targeted for degradation by dTAG-13 prompted us to confidently proceed with the quantitative phospho-proteomic approach to determine putative PPP2CA substrates. Quantitative phospho-proteomic analysis in DMSO- and dTAG-13-treated ^dTAG/dTAG^PPP2CA HEK293 cell extracts identified a total of 39,103 phospho-peptides belonging to 5,829 proteins (full list available in Table S2). Of these, 2,651 phospho-peptides corresponding to 1,149 proteins showed a significant increase in abundance of >2-fold in dTAG-13-treated cells compared to DMSO-treated controls, while 6,280 phospho-peptides corresponding to 2,204 proteins showed a significant increase in abundance of >1.5-fold (Fig. 2C). These phospho-peptides denote potential PPP2CA substrates in these cells under the experimental conditions employed. In contrast, only 16 phospho-peptides belonging to 11 proteins were found to be significantly reduced in abundance by >1.5-fold in dTAG-13-treated cells compared to DMSO-treated controls, which could imply indirect phosphorylation of targets that is dependent on PPP2CA activity or abundance.

**Figure 2.**
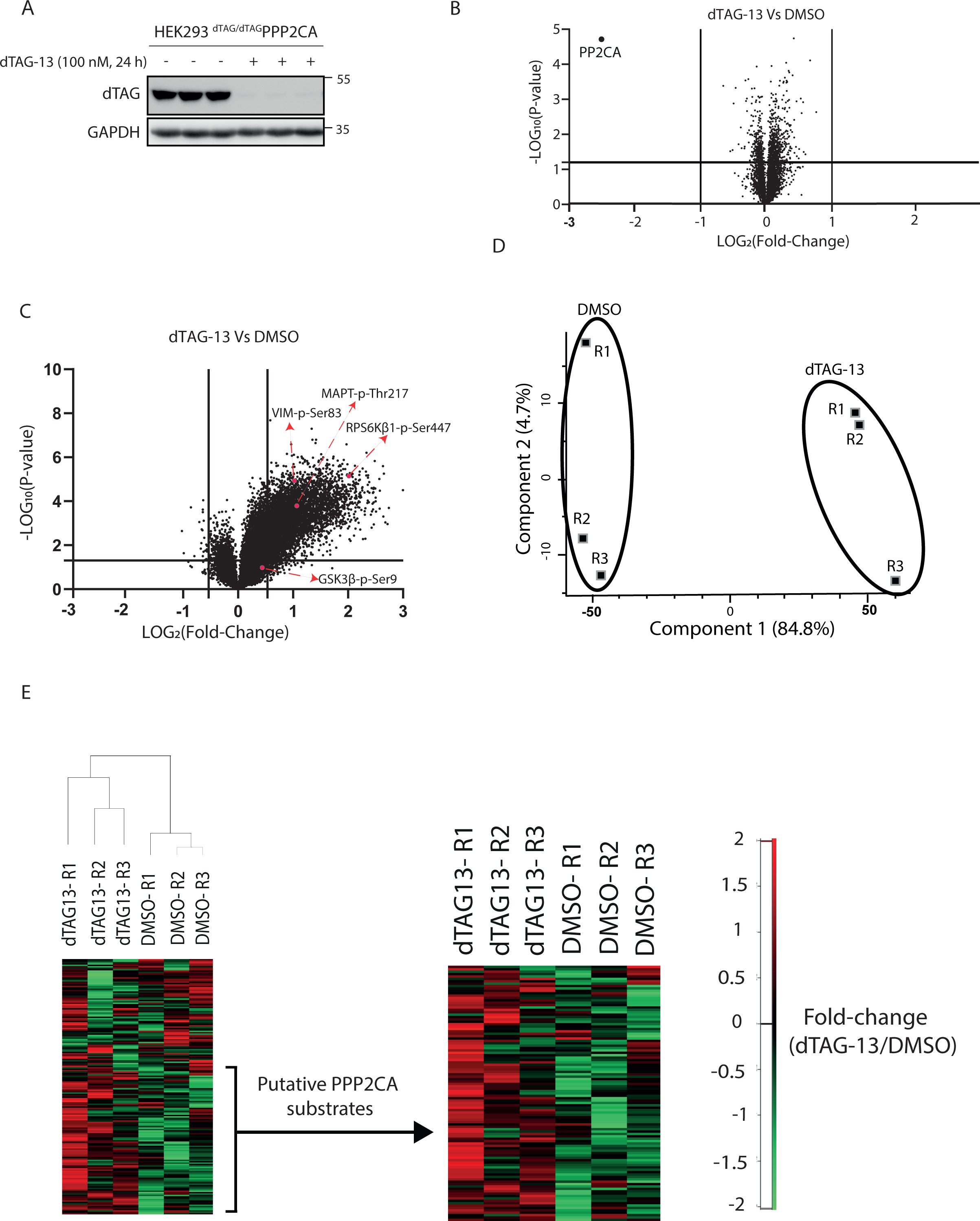
Global proteomic and phospho-proteomic analysis upon PPP2CA degradation. **A.** Extracts (20 μg protein) from ^dTAG/dTAG^PPP2CA HEK293 cells treated for 24 h with DMSO or dTAG-13 (100 nM) (3 independent treatments per condition) prior to lysis were resolved by SDS-PAGE and transferred to nitrocellulose membranes, which were analysed by immunoblotting with the indicated antibodies. **B.** Extract samples from (A) were analysed using the TMT-based proteomics workflow, as described in Fig. S4. Volcano plot showing quantitative changes in the identified proteins. Data plotted represent log_2_ of the fold change in protein abundance in dTAG-13-treated extracts normalised to DMSO-treated controls against −log_10_ of the P value for each identified protein. Under these conditions, the only protein that was significantly degraded (FC <0.5) upon dTAG-13 treatment was dTAG-PPP2CA. **C.** Volcano plot showing global phospho-proteome alteration in dTAG-13-treated ^dTAG/dTAG^PPP2CA HEK293 cells compared to DMSO-treated controls. Data plotted represent log_2_ of the fold change of phospho-peptides identified in dTAG-13-treated extracts normalised to DMSO-treated controls against −log_10_ of the P value for each phospho-peptide. The positions of some phospho-proteins of interest are indicated by arrows. **D.** Principal component analysis of global quantitative phospho-proteomics data across the three biological replicates (R1-3) for each condition. **E.** Unsupervised clustering of altered phospho-peptides identified in dTAG-13- and DMSO-treated ^dTAG/dTAG^PPP2CA HEK293 cells. ANOVA was used to identify significantly altered phospho-peptides upon dTAG-PPP2CA degradation. Abundance values of differentially phosphorylated peptides are represented in a heatmap format, with green representing low abundance and red representing high abundance. The sliding-scale for relative phospho-protein abundance is included. Phospho-peptides whose abundance significantly increased upon dTAG-13-mediated dTAG-PPP2CA degradation, indicating them as putative PPP2CA substrates, are shown with increased magnification.

To ensure concordance in identified phospho-peptides between the biological replicates for both DMSO- and dTAG-13-treated cells, a principal component analysis (PCA) was conducted using Perseus (Fig. 2D). Indeed, PCA revealed that individual replicates from each group cluster together, with a clear separation between DMSO and dTAG-13 groups (Fig. 2D). A heat map was generated through hierarchical clustering of the identified phospho-peptides and potential PPP2CA substrates, which denote phospho-peptide enrichment in dTAG-13-treated replicates relative to DMSO-treated replicates, are indicated (Fig. 2E). We took a selection of some interesting identified phospho-peptides and analysed the significance in their phosphorylation upon dTAG-PPP2CA degradation using violin plots (Fig. S5). These data clearly demonstrate a significant enrichment of these phospho-peptides upon dTAG-PPP2CA degradation, suggesting them to be putative PPP2CA substrates.

### *In silico* analysis of identified phospho-peptides and proteins as potential PPP2CA targets

To elucidate whether the phospho-peptides enriched upon dTAG-PPP2CA degradation conform to a consensus dephosphorylation motif, a multiple sequence alignment was conducted for phospho-peptides that were enriched >2-fold in dTAG-13-treated cells over DMSO-treated controls, by using 16-mer peptides with the identified phospho-Ser/Thr residue placed in the middle (Schneider and Stephens, 1990, Crooks et al., 2004). Consistent with the reported role of PPP2CA in dephosphorylating p-Ser/p-Thr residues, all enriched phospho-events were observed on Ser and Thr residues, with a majority (70%) observed on Ser residues (Fig. 3A), which contrasts the intrinsic phospho-Thr preference that was reported for recombinant PPP2CA substrates *in vitro* (Hoermann et al., 2020) and highlights the likely impact of the *in cellulo* context represented by our phospho-proteomics. In almost 50% of the identified phospho-peptides, enrichment was observed for peptides possessing a Pro residue at the +1 position. This is consistent with previous studies that have reported some PP2A-mediated dephosphorylation of p-Ser/Thr residues that are phosphorylated by Pro-directed kinases, such as sites on microtubule-associated protein tau (Qian et al., 2010, Liu et al., 2005).

**Figure 3.**
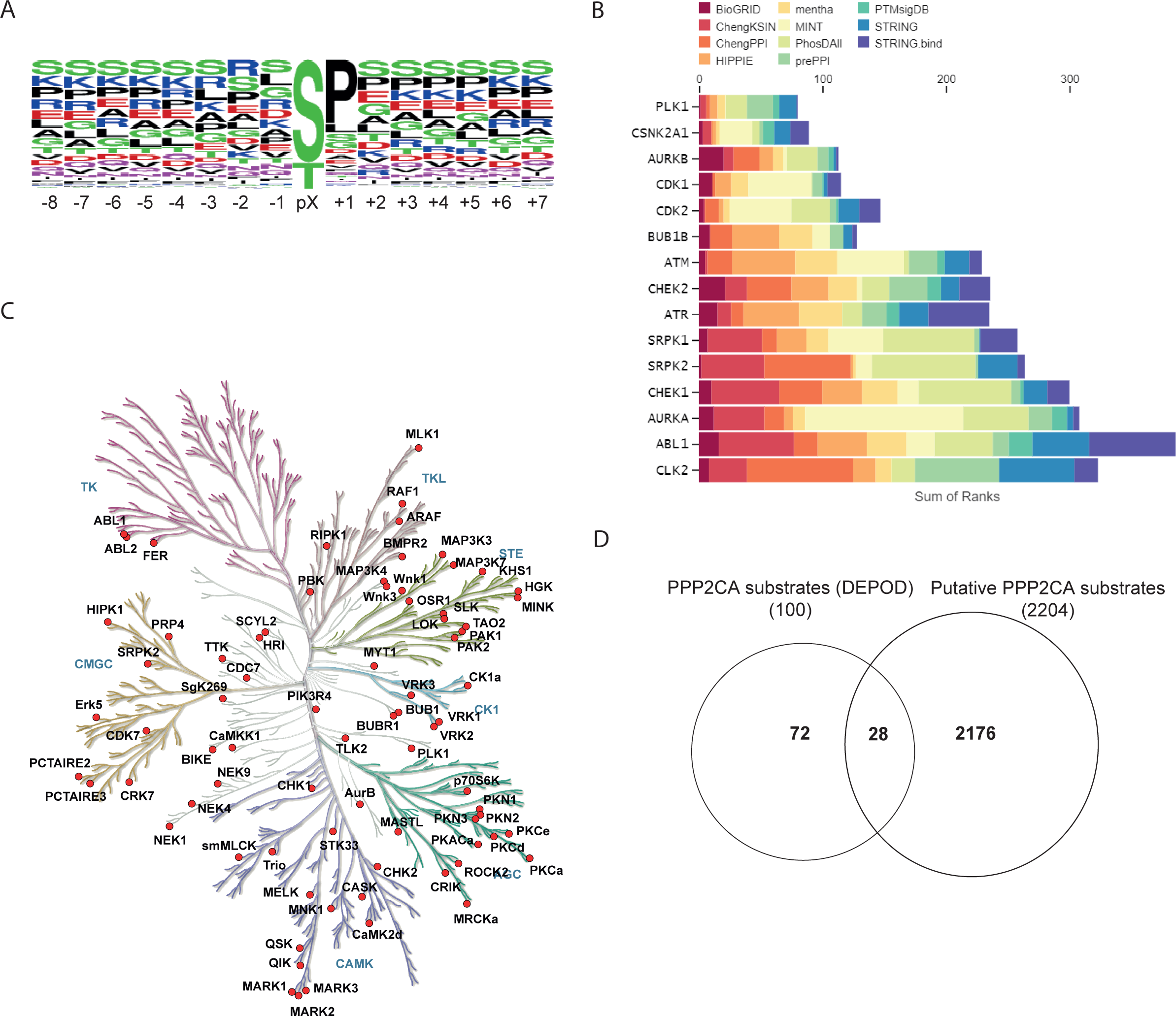
Deconvolution of the significantly enriched phospho-peptides upon dTAG-PPP2CA degradation. **A.** Motif analysis from the significantly enriched phospho-peptides upon dTAG-PPP2CA degradation. **B.** Bar chart representing the MeanRank visualization from the Kinase Enrichment analysis based on phospho-proteins enriched upon dTAG-PPP2CA degradation. The top 15 kinases are plotted against the integrated ranking of the predicted kinases known to phosphorylate the identified phospho-proteins across different libraries, based on MeanRank score. The bar is colour-coded to reflect each library used for data analysis. **C.** Protein kinases for which phospho-peptides were significantly enriched upon dTAG-PPP2CA degradation are indicated on the kinome map, which was generated using the KinMap tool. These kinases are potentially regulated by PPP2CA. **D.** Venn diagram comparison between the identified putative PPP2CA substrates from our datasets and the reported PP2A substrates from the DEPOD database.

At the −1 position, enrichment of Ser, Leu, Gly, Arg, Asp and Lys residues was observed, indicating a tolerance for any charged or hydrophobic residues immediately N-ter of the phospho-site. Further upstream and downstream of the phospho-site, enrichment was seen for Ser, Lys, Arg, Pro, Glu and Thr residues, suggesting that PPP2CA dephosphorylation weakly favours upstream and downstream residues that possess electrically charged or polar side chains. Enrichment of Ser and Thr residues both upstream and downstream of the phospho-site may imply that PPP2CA dephosphorylates clusters of multiple phospho-Ser/Thr residues. Indeed, amongst the phospho-peptides enriched upon dTAG-PPP2CA degradation, ∼4000 were identified as mono-phosphorylated, ∼1500 as di-phosphorylated and ∼300 as tri-phosphorylated (Fig. S6A). Having explored the consensus motif of the PPP2CA-regulated phospho-sites, we were interested to explore the prevalence of known PP2A-B56α SLiM motif LSPIxE in our identified putative PPP2CA substrates. For this, FIMO (Find individual Motif Occurrences) was employed to scan the human proteome for the motif LSPIxE, which identified 897 unique proteins as containing this SLiM. Of these, 99 were identified by our phospho-proteomic analysis (Fig. S6B), suggesting these may be PP2A-B56α substrates. Absence of this SLiM from other identified putative PPP2CA substrates potentially indicates other modes of substrate recruitment. In a similar vein, failure to identify other LSPIxE-motif containing proteins as potential PPP2CA substrates in our study could imply that these proteins are either not substrates of PPP2CA or the absence of an appropriate biological context in our study may account for lack of their identification.

We undertook an *in silico* analysis to identify the upstream kinases whose putative substrates are overrepresented in our identified putative PPP2CA substrate phospho-proteins. Through the Kinase Enrichment Analysis 3 tool, we found Ser/Thr-protein kinase PLK1, casein kinase 2α (CK2α), aurora kinase A and B (AURKA/B), cyclin-dependent kinase 1 and 2 (CDK1 and CDK2), BUB1B/BUBR1, ataxia-telangiectasia mutated (ATM), checkpoint kinase 1 and 2 (CHEK1/2), ataxia telangiectasia and Rad3-related protein (ATR), Ser/Arg-rich splicing factor protein kinase 1 and 2 (SRPK1/2), Tyr protein kinase ABL1 and dual specificity CDC-like kinase 2 (CLK2) as among the top kinases known to regulate the phospho-proteins that we identified as putative PPP2CA substrates (Fig. 3B). We also mapped all protein kinases within the human kinome tree using KinMap to reveal the 50 kinases whose phosphorylation at certain residues was enriched upon dTAG-PPP2CA degradation (Fig. 3C), suggesting PPP2CA activity might regulate these kinases, which are involved in cell processes such as mitosis, proliferation and gene expression (Glover et al., 1995, Wang, 2014, Fisher, 2019). Additionally, we compared the identified putative PPP2CA substrates against the reported PPP2CA substrates on the Human Dephosphorylation Database, DEPOD (Damle and Kohn, 2019). Of the 2,204 most significantly enriched phospho-proteins from our study, only 28 were reported by the DEPOD as PP2A substrates (Fig. 3D), indicating that our study has provided a step change in the number of phospho-proteins reported as potential PPP2CA substrates. Furthermore, we compared the proteins our study identified as putative PPP2CA substrates with those from previous studies, and our study identified 262 out of 515 phospho-proteins identified as *in vitro* PPP2CA substrates (Hoermann et al., 2020) (Fig. S6C) and 345 out of 522 phospho-proteins PPP2R1A substrates (Kauko et al., 2020) (Fig. S6D). In each case, our approach identified >1800 unique phospho-proteins as putative PPP2CA substrates. Encouragingly, when comparing to the DEPOD database, Enrichr listed PPP2CA as the phosphatase predicted to be the most likely to dephosphorylate the phospho-proteins we identified in our study as putative PPP2CA substrates (Fig. S6E).

### *In silico* prediction of biological roles of the phospho-proteins identified as putative PPP2CA substrates

From the identified phospho-proteins that were enriched upon dTAG-PPP2CA degradation, we explored *in silico* analysis of their reported involvement in cell signalling pathways, biological processes, molecular function, protein domain architecture, subcellular distribution, disease associations and protein-protein interactions (Fig. 4A-F and Fig. S7). Analyses were conducted using the Database for Annotation, Visualization and Integrated Discovery (DAVID), Enrichr, FunRich and STRING (Huang da et al., 2009, Sherman et al., 2022, Chen et al., 2013, Kuleshov et al., 2016, Xie et al., 2021, Pathan et al., 2015, Snel et al., 2000) and the full lists of results can be found in Table S3. Kyoto Encyclopedia of Genes and Genomes (KEGG) pathway analysis revealed that a significantly high percentage of the identified phospho-proteins are known to be involved in spliceosome function, the cell cycle, RNA transport and surveillance, ubiquitin-mediated proteolysis, endocytosis and DNA replication (Fig. 4A). The identified phospho-proteins were also significantly implicated in biological processes including regulation of transcription by RNA polymerase II, DNA transcription, chromatin organisation and mRNA splicing (Fig. 4B). These observations were supported by the reported molecular functions of the identified phospho-proteins, which showed a significantly high percentage to be involved in binding RNA, cadherin, mRNA, DNA, microtubules and tubulin (Fig. 4C). When we performed a network analysis of the phospho-proteins, major functional groups identified showed interaction of groups involved in the spliceosome, RNA transport, DNA replication, DNA repair and cell cycle (Fig. S7), supporting the KEGG pathway, biological processes and molecular function associations above. Domain architecture analysis of the identified phospho-proteins revealed that 43% contained coiled-coil domains, which are prominent in transcriptional regulation (Fig. 4D). Other domains that showed a significant presence in the identified phospho-proteins included the RNA recognition motif (RRM) (5.6%), plant homeodomain (PHD) (2.8%), bromo domain (BROMO) (1.5%), helicase superfamily c-terminal domain (HELICc) (2.4%), and the forkhead-associated (FHA) domain (FHA) (1.1%). Analysis of the subcellular distribution of the identified phospho-proteins indicated presence in many subcellular compartments, including the nucleus, membrane- and non-membrane-bound organelles and microtubule cytoskeleton (Fig. 4E). Disease associations of the identified putative PPP2CA substrates showed involvement in micrognathism, developmental delay, microcephaly, vesicoureteral reflux, mental retardation, congenital epicanthus, breast cancer and neurodevelopmental disorders (Fig. 4F).

**Figure 4.**
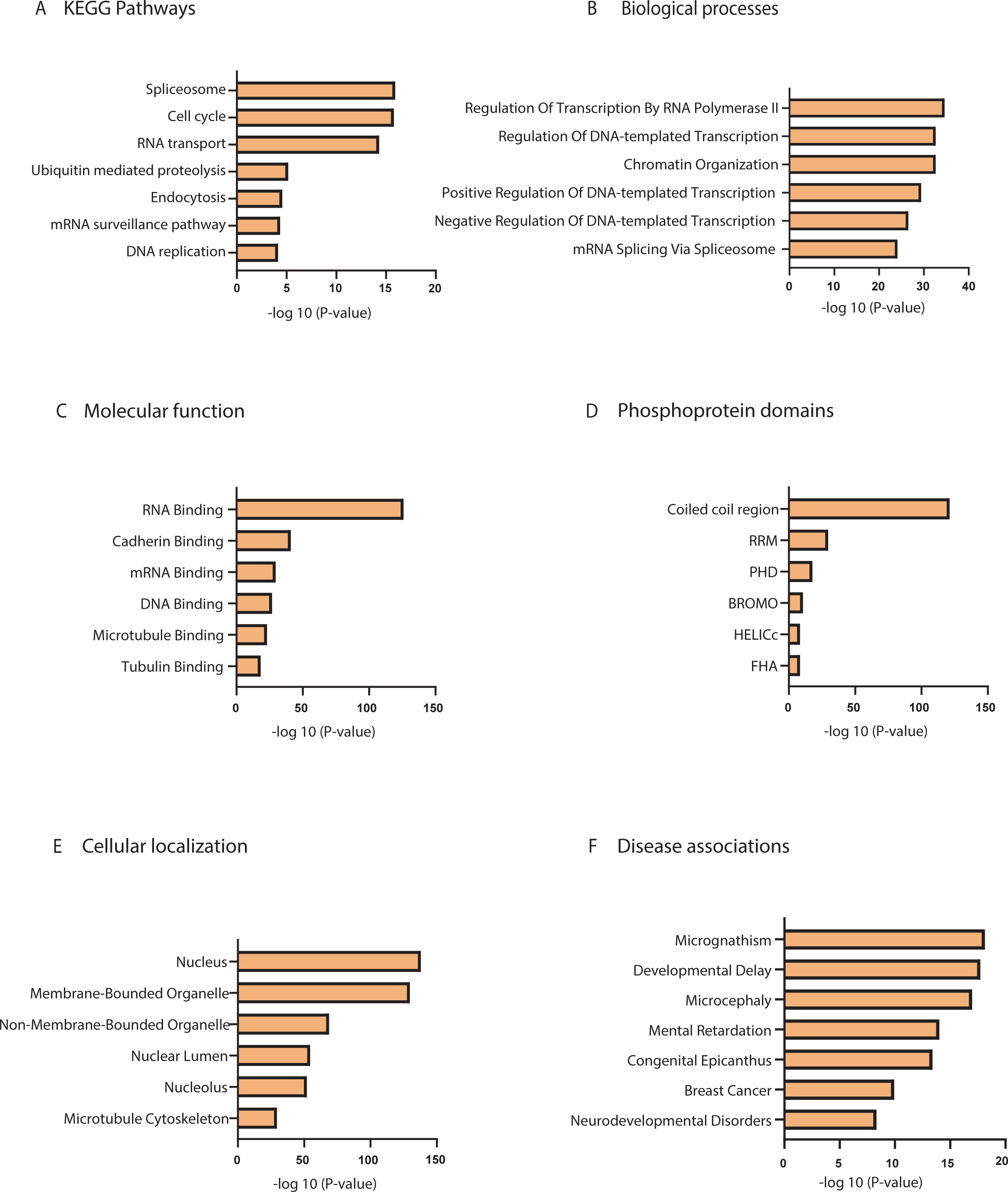
Gene Ontology (GO) analysis of proteins identified as putative PPP2CA substrates. **A.** Involvement of the identified putative PPP2CA substrates in KEGG (Kyoto Encyclopedia of Genes and Genomes) pathways. **B.** Involvement of the identified putative PPP2CA substrates in different biological processes. **C.** Known and predicted molecular functions of the identified putative PPP2CA substrates **D.** Domain architecture of proteins identified as putative PPP2CA substrates. **E.** Known subcellular distribution of the identified putative PPP2CA substrates. **F.** Disease associations of the identified putative PPP2CA substrates For A-D, the data is represented as bar charts indicating the p-value for associations of the identified putative PPP2CA substrates to given biological pathways (A), cellular processes (B), molecular functions (C), domain architectures (D), subcellular distribution (E) and diseases (F). These values were generated using gene ontology (GO) analysis and a selection of only the top classes have been included in the bar charts.

### Validation of PPP2CA phospho-proteomic data by immunoblotting

To validate some of the phospho-proteins identified by phospho-proteomics as potential PPP2CA targets, WT and ^dTAG/dTAG^PPP2CA HEK293 cells were treated with DMSO, MLN4924 (1 µM), dTAG-13 (100 nM) or co-treatment of dTAG-13 and MLN4924 for 24 h prior to lysis. Immunoblotting with anti-dTAG antibody confirmed the degradation of dTAG-PPP2CA in ^dTAG/dTAG^PPP2CA HEK293 cells treated with dTAG-13 in comparison to DMSO (Fig. 5A). Treatment of ^dTAG/dTAG^PPP2CA HEK293 cells with MLN4924 alone had no effect on dTAG-PPP2CA abundance, while co-treatment of cells with MLN4924 and dTAG-13 rescued the dTAG-PPP2CA degradation caused by dTAG-13 (Fig. 5A). Again, MLN4924 treatment resulted in inhibition of cullin NEDDylation and activation, evident by the band collapse of CUL2 in MLN4924-treated samples (Fig. 5A). No changes in the abundance of endogenous PPP2CA/B in WT cells and PPP2CB in ^dTAG/dTAG^PPP2CA HEK293 cells were apparent when cells were treated with DMSO, MLN4294, dTAG-13 or dTAG-13+MLN4924 (Fig. 5A). Under these conditions, we probed these extracts with phospho-specific antibodies against some of the phospho-proteins that we identified as potential PPP2CA targets from the phospho-proteomic analysis.

**Figure 5.**
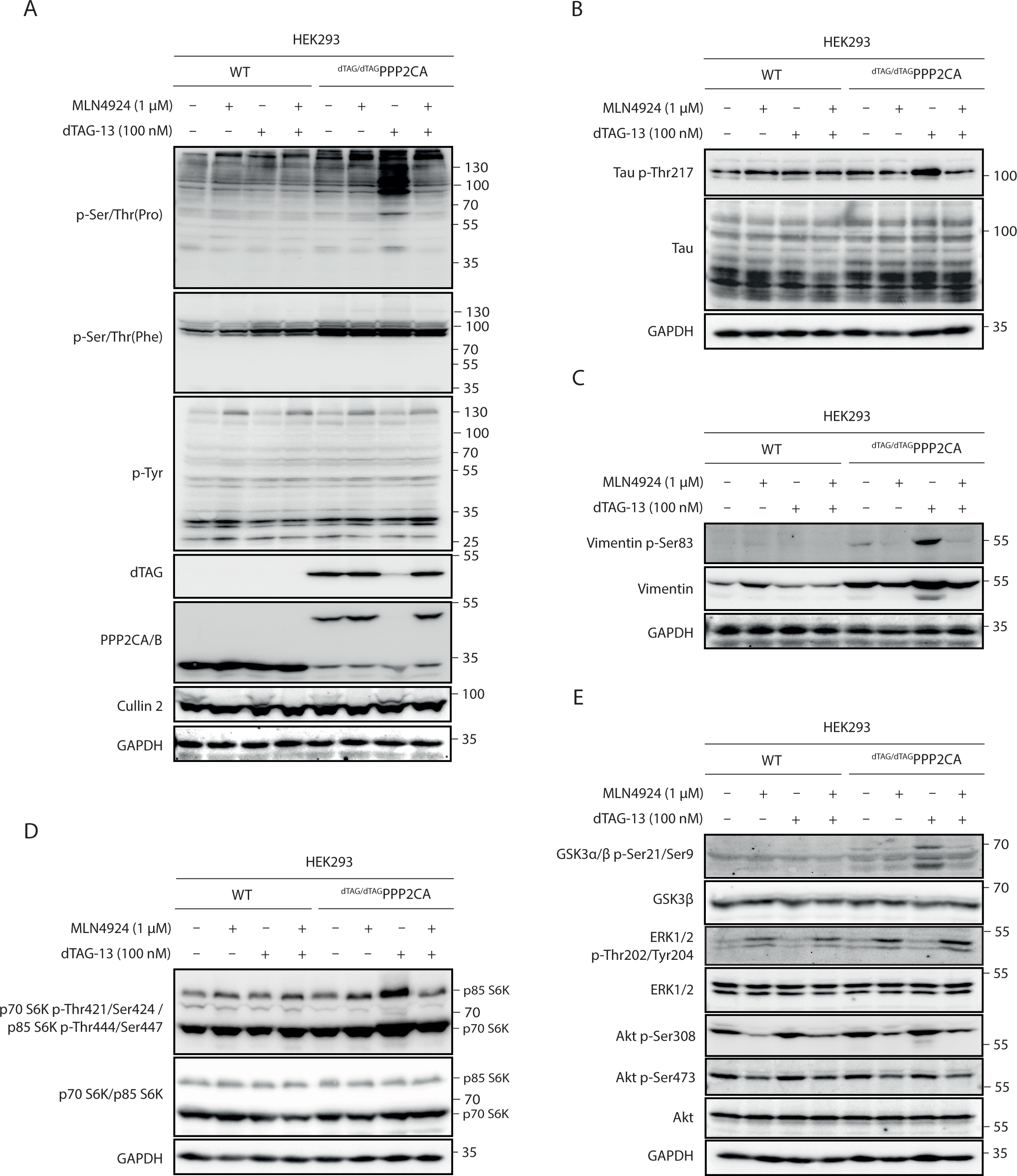
Immunoblot validation of putative PPP2CA substrates. **A-D.** Extracts (20μg protein) from WT and ^dTAG/dTAG^PPP2CA HEK293 cells treated with DMSO, dTAG-13 (100 nM), MLN4924 (1000 nM) or dTAG-13 and MLN4924 for 24 h prior to lysis were resolved by SDS-PAGE and transferred to nitrocellulose membranes, which were analysed by immunoblotting with the indicated antibodies. Data representative of 3 independent immunoblots.

Just as phospho-peptides that conform to a phospho-Ser/Thr-Pro motif were overwhelmingly enriched upon dTAG-PPP2CA degradation from phospho-proteomic analysis (Fig. 3A), immunoblotting with an anti-phospho-Ser/Thr-Pro motif antibody corroborated these observations (Fig. 5A). Levels of phospho-Ser/Thr-Pro were greatly enhanced upon degradation of dTAG-PPP2CA with dTAG-13 in ^dTAG/dTAG^PPP2CA HEK293 cells but not in WT HEK293 cells, in comparison to corresponding DMSO-treated controls (Fig. 5A). MLN4924 treatment alone in both cells caused little difference to phospho-Ser/Thr-Pro signals relative to DMSO-treated controls. However, in ^dTAG/dTAG^PPP2CA HEK293 cells treated with dTAG-13, co-treatment with MLN4924, which prevented the degradation of dTAG-PPP2CA, caused a reduction in phospho-Ser/Thr-Pro signals to similar levels observed with DMSO or MLN4924 treatment alone. In contrast to the phospho-Ser/Thr-Pro motif, there was no discernible change in either phospho-Ser/Thr-Phe or phospho-Tyr levels across the samples (Fig. 5A), which is consistent with the *in silico* analysis of the phospho-proteomic data, which showed no enrichment of these motifs among the identified phospho-peptides (Fig. 3A).

We also sought to validate individual potential PPP2CA substrates identified from phospho-proteomic analysis. Despite a limited availability of phospho-specific antibodies raised against the particular phospho-sites elucidated by phospho-proteomic analysis, we were able to validate a small number of these potential PPP2CA substrates. Firstly, our phospho-proteomic analysis identified a pThr534 peptide from tau to be significantly enriched by 1.9-fold in dTAG-13-treated ^dTAG/dTAG^PPP2CA HEK293 cells compared to DMSO-treated controls (Table S2). By immunoblotting with an anti-pThr217 (also known as p-Thr534) tau antibody, we observed a clear increase in pThr217 abundance in dTAG-13-treated ^dTAG/dTAG^PPP2CA HEK293 cells in comparison to DMSO-treated controls (Fig. 5B). In contrast, treating WT HEK293 cells with dTAG-13 did not yield an increased pThr217 tau abundance in comparison to DMSO treatment (Fig. 5B. The pThr217 signal was only detected on a tau species of ∼100 kDa, corresponding to Big-tau, which contains a large additional exon termed exon 4a (Fischer and Baas, 2020). Total tau levels, regardless of the isoforms, remained similar across all treatment conditions. Co-treatment of ^dTAG/dTAG^PPP2CA HEK293 cells with dTAG-13 and MLN4924, which rescued dTAG-PPP2CA degradation, reduced the pThr217 tau abundance to levels seen with DMSO treatment or MLN4924 treatment alone (Fig. 5B). PP2A has previously been reported to dephosphorylate tau at multiple residues, including Thr205, Thr212, Ser214 and Ser262 (Qian et al., 2010). Of these, our proteomic screen significantly identified Thr212 and Ser214 from Big-tau (Thr529 and Ser531) with fold changes of 1.9 and 1.2, respectively (Table S2).

Our phospho-proteomic analysis also revealed a significant 1.8-fold change of a phospho-peptide containing pSer83 of vimentin, which was previously reported to be hyperphosphorylated upon PP2A inhibition (Table S2) (Turowski et al., 1999). By immunoblot with anti-vimentin pSer83 antibody, we observed a substantial increase in pSer83-vimentin levels in dTAG-13-treated ^dTAG/dTAG^PPP2CA HEK293 cells in comparison to DMSO treatment, while a slight increase in total vimentin levels and an extra lower molecular weight species were also apparent (Fig. 5C). Hyperphosphorylation of vimentin is reported to cause disassembly of vimentin filaments into bundles around the nucleus (Turowski et al., 1999), which may be related to the change in abundance and the appearance of an additional vimentin species that we observed here. No substantial changes in pSer83-vimentin levels were evident in WT HEK293 cells with any of the treatments.

A phospho-peptide containing p-Ser447 of p85 S6K was also identified in our phospho-proteomic analysis and experienced a significant 3.6-fold change in abundance in dTAG-13-treated ^dTAG/dTAG^PPP2CA HEK293 cells in comparison to DMSO-treated controls (Table S2). p85 S6K and p70 S6K are two protein isoforms formed through alternative splicing of the ribosomal protein S6 kinase β1 (RPS6Kβ1) gene, with p85 S6K containing an extra 23 amino acids at the N-terminus in comparison to the p70 S6K isoform (Reinhard et al., 1992). These S6K isoforms are Ser/Thr protein kinases that respond to mammalian target of rapamycin (mTOR) signalling and act downstream of phosphoinositide-dependent protein kinase 1 (PDPK1) to phosphorylate the S6 ribosomal protein, resulting in an increase in protein synthesis and cell proliferation (Chung et al., 1992, Chung et al., 1994). RPS6Kβ1 has been reported as a PP2A substrate previously (Ballou et al., 1988, Peterson et al., 1999, Petritsch et al., 2000). When immunoblotting with an anti-phospho-S6K antibody, which recognises both p-Thr444/Ser447 p85 S6K and p-Thr421/Ser424 p70 S6K, a clear increase in signal intensity was observed for p85 S6K as well as p70 S6K in dTAG-13-treated ^dTAG/dTAG^PPP2CA HEK293 cells in comparison to DMSO treatment and dTAG-13-treated WT HEK293 cells (Fig. 5D). Interestingly, the levels of phospho-S6K were seen to increase with MLN4924 treatment, in both WT and ^dTAG/dTAG^PPP2CA HEK293 cells, suggesting that inhibition of cullin activity may impact the stability of phospho-S6K, or the phosphorylation of S6K at these residues (Fig. 5D).

Finally, a phospho-peptide containing p-Ser9 and corresponding to reported PP2A-regulated protein GSK3β was identified with a fold change of 1.4 in dTAG-13-treated cells in comparison to DMSO-treated controls, but with a p-value >0.05 (Table S2) (Cross et al., 1995, Shaw et al., 1997, Mitra et al., 2012, Wang et al., 2015). By immunoblot, the levels of p-Ser21/9 GSK3α/β were substantially increased in ^dTAG/dTAG^PPP2CA HEK293 cells treated with dTAG-13 over DMSO, while no changes were observed with dTAG-13 or DMSO treatment in WT HEK293 cells (Fig. 5E). Co-treatment of dTAG-13-treated ^dTAG/dTAG^PPP2CA HEK293 cells with MLN4924, which rescues dTAG-PPP2CA degradation, caused a return in the levels of p-Ser21/9 GSK3α/β to those seen with DMSO treatment, while MLN4924 treatment alone had no effect (Fig. 5E). These data suggest that perhaps, due to the vast number of changing phospho-peptides in response to dTAG-PPP2CA degradation and subsequent normalisation, the phospho-peptide containing p-Ser9 GSK3β was not detected to be significantly enriched in our phospho-proteomic analysis. We also tested some other proteins that have been reported to be PP2A substrates or markers within PP2A-regulated pathways, including Akt and extracellular signal-regulated kinase 1/2 (ERK1/2) (Kuo et al., 2008, Mao et al., 2005, Letourneux et al., 2006), which did not appear in our phospho-proteomic screen (Fig. 5E). Only a modest increase in phosphorylation was observed for Akt p-Thr308 following dTAG-PPP2CA degradation (Fig. 5E). This is likely due to the fact that our experiments were performed in conditions where ERK1/2 and Akt signalling pathways were not stimulated with growth factors to induce high levels of phosphorylation of these target proteins.

## Discussion

In this study we combined CRISPR/Cas9 genome editing, PROTAC-mediated targeted protein degradation and unbiased phospho-proteomics to identify over two thousand proteins as putative substrates of PPP2CA, the major catalytic subunit of the PP2A holoenzyme complex. Some among these have been reported as PP2A substrates, but a vast majority are novel putative substrates, thus offering great potential to better understand substrate- and pathway-level phospho-regulation within cell signalling. Quantitative total proteome analysis upon targeted dTAG-PPP2CA degradation over 24 h revealed that PPP2CA was the only protein whose abundance was significantly altered, confirming the exquisitely selective nature of PROTAC-mediated POI degradation and also suggesting that inhibition of PPP2CA activity does not appear to impact the stability of its substrates. Despite the identification of >6000 phospho-peptides as putative PPP2CA substrates upon its degradation over 24 h, surprisingly this did not cause substantial cytotoxicity over this time period. More predictably, sustained PPP2CA degradation over 48 h and 72 h caused a significant inhibition of cell proliferation.

Given that we conducted the phospho-proteome analysis in ^dTAG/dTAG^PPP2CA HEK293 cells under cell culture conditions, a limitation of our study is the lack of biological or cell signalling context. However, the phospho-proteomics workflow and the cells we employed here can be applied to any cell signalling context for dissecting PPP2CA substrates in specific contexts. As with any phospho-proteomic study, the detection of putative substrates relies on the design and sensitivity of the approach, and a lack of detection of putative substrates potentially means that the number of PP2A substrates that we have identified is an underestimate of the true number of substrates. While our high-throughput phospho-proteomics method successfully identified nearly 40,000 unique phospho-peptides, a significant portion of crucial regulatory phosphorylation events has proven elusive. This challenge arises because many sites are inaccessible or challenging to detect when subjected to trypsin digestion, which hinders MS-based investigations. Protein phosphorylation, particularly phosphorylation of multiple residues in close clusters, affects both proteolytic cleavage and ionisation, thereby affecting detection by mass-spectrometry. Utilising multiple proteases in the phospho-proteomic analysis could be advantageous as it could improve throughput. Some of these issues could potentially account for other reported PPP2CA substrates not being significantly identified in our study, such as phospho-GSK3β, which we were able to validate by immunoblotting. Importantly, for some of the phospho-peptides that we identified as putative PPP2CA substrates by mass spectrometry, we were able to validate them by immunoblotting, suggesting our data to be robust. Given that the field of protein phosphatase research has lagged that of protein kinases, our approach has the potential to expedite the pace of research into phosphatases by identifying substrates in different biological settings, although care must be taken to ensure that the introduction of a degron tag does not compromise the function of the phosphatase in the first place.

*In silico* analysis of the phospho-peptide sequences enriched upon dTAG-PPP2CA degradation identified proline-directed p-Ser/p-Thr residues as key targets of PPP2CA. Other motif elements that we uncovered might provide insights into the residues upstream and downstream of the phospho-site that are or are not tolerated for PPP2CA-mediated dephosphorylation. Analysis of putative PP2A substrate proteins revealed their involvement in many key biological pathways, including spliceosome function, the cell cycle, RNA transport and surveillance, ubiquitin-mediated proteolysis and DNA repair, which is consistent with the reported pleiotropic roles of PP2A holoenzyme complexes (Wlodarchak and Xing, 2016, Reynhout and Janssens, 2019). Future studies will need to establish whether the PPP2CA-regulated dephosphorylation of individual proteins correlates with their functions in these biological processes and whether PPP2CA is directly or indirectly responsible for substrate dephosphorylation. Similarly, many enzymes, such as protein kinases and E3 ubiquitin ligases, were also identified as putative PPP2CA substrates, elucidating potential crosstalk between key regulators of cells signalling. For most of the identified phospho-sites, it is not known how phosphorylation affects the activities or behaviour of these enzymes. We observed a significant association of putative PPP2CA substrates with breast cancer as well as numerous developmental disorders, such as micrognathism, developmental delay, microcephaly, mental retardation, congenital epicanthus and neurodevelopmental disorders. Some putative PPP2CA substrates we identified that have previously been linked to neurodevelopmental disorders include RING finger protein 12 (RNF12), OTU deubiquitinase 5 (OTUD5), methyl CpG binding protein 2 (MECP2), microcephalin 1 (MCPH1), fragile X mental retardation protein (FMRP), Ser/Arg-rich splicing factor protein kinase 2 (SRPK2) and budding uninhibited by benzimidazoles 1 (BUB1) (Bustos et al., 2020, Beck et al., 2021, Stefanelli et al., 2016, Meyer et al., 2019, Sidorov et al., 2013, Carvalhal et al., 2022). RNF12 (also known as RLIM) is an E3 ligase that promotes the ubiquitin-mediated degradation of the transcription factor reduced expression 1 (REX1, also known as Zfp42), thus preventing transcription of neural genes (Barakat et al., 2011, Shin et al., 2010). RNF12-dependent ubiquitination of REX1 was found to be stimulated following RNF12 phosphorylation by the SRPK (Ser/Arg-rich splicing factor (SRSF) protein kinase) kinase family (Bustos et al., 2020). Interestingly, we observed enrichment of SRPK2-regulated phospho-sites upon dTAG-PPP2CA degradation, and even some phospho-sites on SRPK2 itself, potentially offering some insight into the association of hyperphosphorylated hits with neurodevelopmental disorders. Previously, de novo mutations in PPP2CA, and another PP2A subunit, PPP2R5D, have been identified in patients with intellectual disability and developmental delay (Mirzaa et al., 1993, Reynhout et al., 2019). Furthermore, mice with conditional loss of PPP2CA in the central nervous system were reported to exhibit severe microcephaly, cortical atrophy and intellectual learning and memory defects (Liu et al., 2018, Reynhout and Janssens, 2019), supporting an important role for PPP2CA in the context of development.

Our study demonstrates that targeted protein degradation and subsequent global phospho- and total-proteomic analysis is a robust approach to interrogate the substrates of protein phosphatases. PPP2CA is one of the catalytic subunits of the PP2A holoenzyme complex, with the other one being PPP2CB, which is 97% identical (Khew-Goodall et al., 1991). Since our data suggests that PPP2CB was still expressed in ^dTAG/dTAG^PPP2CA HEK293 cells, it appears that PPP2CB was unable to fully compensate for the loss of PPP2CA in these cells, potentially suggesting unique roles for these highly similar proteins. It would be interesting to identify PPP2CB substrates using a similar approach and to combine PPP2CA and PPP2CB degradation together, to uncover unique and common PPP2CA and PPP2CB substrates, thus providing a full understanding of the extent of dephosphorylation conducted by PP2A holoenzyme complexes in cells.

## Materials and Methods

### Materials Availability

All constructs used in this study are available to request from the MRC PPU Reagents & Services webpage (http://mrcppureagents.dundee.ac.uk) with the unique identifier (DU) numbers providing direct links to the cloning strategies and sequence details. All constructs were sequence-verified by the DNA Sequencing Service, University of Dundee (http://www.dnaseq.co.uk).

## Experimental models and subject details

### Cell Lines

Aseptic technique that meets biological safety requirements was used for all procedures. HEK293 cells are immortalised human embryonic kidney cells derived from a female foetus. HEK293 cells were cultured in DMEM (Life Technologies) supplemented with 10% (v/v) foetal bovine serum (FBS, Thermo Fisher Scientific), 2 mM L-glutamine (Lonza), 100 U/ml penicillin (Lonza) and 0.1 mg/ml streptomycin (Lonza). Cells were maintained at 37°C with 5% CO_2_ in a water-saturated incubator. For passaging, trypsin/EDTA was used at 37°C to detach cells.

## Methods

### Generation of cell lines using CRISPR/Cas9

The CRISPR/Cas9 genome editing system (Ran et al., 2013) was used to generate HEK293 *PPP2CA* homozygous N-terminal dTAG KI (^dTAG/dTAG^PPP2CA) cells. HEK293 wild type cells were transfected with vectors encoding a guide RNA (gRNA) targeting the PPP2CA exon 1 locus (DU69331, pX459 puromycin Cas9^D10A^ PPP2CA) (1 µg) and donor (DU69361, pMA PPP2CA Nter GFP IRES2 FKBP12^F36V^) (3 µg), as well as polyethylenimine (PEI). 16 hr post-transfection, selection with 1 μg/mL puromycin (Sigma-Aldrich) was carried out for 48 hr. The transfection process was replicated (without a further round of selection). Cells were sorted by flow cytometry and single cells were plated in individual wells of 96-well plates. Viable clones were expanded, and integration of dTAG at the target locus was verified by Western blotting, polymerase chain reaction (PCR) amplification and genomic sequencing of the targeted locus.

### Treatment of cells with compounds

The following chemicals were added to cell media using the treatment durations and concentrations indicated in figure legends: dimethylsulphoxide (DMSO) (Sigma-Aldrich), dTAG-13 (MRC PPU Reagents and Services), MLN4924 (MRC PPU Reagents and Services) and MG132 (Merck).

### Cell lysis and immunoprecipitation

Cells were harvested by washing twice with phosphate-buffered saline (PBS) and scraping into ice-cold lysis buffer (50 mM Tris-HCl pH 7.5, 0.27 M sucrose, 150 mM NaCl, 1 mM ethylene glycol-bis(β-aminoethyl ether)-N,N,Nʹ,Nʹ-tetraacetic acid (EGTA), 1 mM ethylenediaminetetraacetic acid (EDTA), 1 mM sodium orthovanadate, 10 mM sodium β-glycerophosphate, 50 mM sodium fluoride, 5 mM sodium pyrophosphate and 1% NP-40) supplemented with 1x cOmplete™ protease inhibitor cocktail (PIC). Lysates were incubated for 10 min on ice before clarification by centrifugation at 17,000 G for 20 min at 4°C. The Bradford assay was used to determine protein concentration and enable normalisation between samples.

### SDS-PAGE and Western blotting

Cell lysates containing equal amounts of protein (20 μg) were resolved by SDS-PAGE and transferred to nitrocellulose membrane. Membranes were blocked in 5% (w/v) non-fat milk (Marvel) in tris-buffered saline, 0.1% Tween 20 (TBS-T) (50 mM Tris–HCl pH 7.5, 150 mM NaCl, 0.2% Tween-20) and incubated overnight at 4°C in 5% (w/v) bovine serum albumin (BSA)/TBS-T or 5% (w/v) milk/TBS-T with the appropriate primary antibodies. Primary antibodies used at indicated dilutions include: anti-Akt (9272, CST, 1:1000), anti-Akt p-Ser308 (4056S, CST, 1:1000), anti-Akt p-Ser473 (4058, CST, 1:1000), anti-Cullin 2 (51-1800, Invitrogen, 1:1000), anti-dTAG (DA179, MRC PPU Reagents and Services, 1 µg/mL), anti-ERK1/2 (9102S, CST, 1:1000), anti-ERK1/2 p-Thr202/Tyr204 (9101S, CST, 1:1000), anti-FKBP12 (ab24373, Abcam, 1:1000), anti-GAPDH-HRP (HRP-60004, ProteinTech, 1:30000), anti-GAPDH (10494-1AP, ProteinTech, 1:30,000), anti-GAPDH (60004-1-Ig, ProteinTech, 1:30,000), anti GSK3-β (9315S, CST, 1:1000), anti GSK3-α/β p-Ser21/Ser9 (9331S, CST, 1:1000), anti-PPP2CA/B (S274B, MRC PPU Reagents & Services, 1 µg/mL), anti-p70 S6 kinase (9202S, CST, 1:1000), anti-p70 S6 kinase p-Thr421/Ser424 (9204, CST, 1:1000), anti-p-Ser/Thr-Phe (9631, CST, 1:1000), anti-p-Ser/Thr-Pro MPM-2 (05-368, Sigma, 1:1000), anti-Tau (S157B, MRC PPU Reagents and Services, 1 µg/mL), anti-Tau p-Thr217 (51625, CST, 1:1000), anti-pTyr (9411, CST, 1:1000), anti-Vimentin (5741S, CST, 1:1000), anti-Vimentin p-Ser83 (3878, CST, 1:1000), anti-Vinculin (13901S, CST, 1:1000).

Membranes were subsequently washed with TBS-T and incubated with horseradish peroxidase (HRP)-or IRDye 800CW-conjugated secondary antibody for 1 hr at room temperature. HRP-coupled secondary antibodies used include: goat anti-rabbit-IgG (7074, CST, 1:5000), rabbit anti-sheep-IgG (31480, Thermo Fisher Scientific, 1:5000), goat anti-mouse-IgG (31430, Thermo Fisher Scientific, 1:5000). IRDye 800CW-coupled secondary antibodies used include: IRDye 800CW Donkey anti-Rabbit IgG (H + L) (926-32213, Licor, 1:5000). After further washing, signal detection was performed using enhanced chemiluminescence (ECL) (Merck) for HRP-conjugated secondaries and ChemiDoc MP System (Bio-Rad). Image Lab (Version 6.0.1) (Bio-Rad) was used to analyse protein bands by densitometry.

### Cell cytotoxicity assay

CellTox Green Assay (Promega, Cat. #G8742) was used to assess the cytotoxicity of dTAG-13-mediated dTAG-PPP2CA degradation in ^dTAG/dTAG^PPP2CA HEK293 cells as well as in control WT HEK293 cells (12 – 48 h, 100 nM dTAG-13 or DMSO control). The fluorescent signal produced by the CellTox Green dye, upon selective binding to the DNA of cells with impaired membrane integrity, is proportional to cytotoxicity. Fluorescence was measured using ex: 480 nm em: 530 nm by a PHERAstar FS plate reader before subtracting blank measurements (made using well containing medium only, no cells) and normalising to DMSO treatment. MG132 treatment (40 µM) was included as a positive control at each time point. Data was analysed using Excel (Microsoft) and GraphPad Prism software (Version 8).

### Cell proliferation assay

The CellTiter 96® AQ_ueous_ One Solution Cell Proliferation Assay (Promega, Cat. #G3580) was used to assess the impact of dTAG-13-mediated dTAG-PPP2CA degradation on cell proliferation in ^dTAG/dTAG^PPP2CA HEK293 cells, as well as in control WT HEK293 cells. Cells were treated with dTAG-13 (100 nM) or DMSO for 24, 48 or 72 h before incubation with the CellTiter 96® AQ_ueous_ One Solution Reagent. The reagent contains a tetrazolium compound [3-(4,5-dimethylthiazol-2-yl)-5-(3-carboxymethoxyphenyl)-2-(4-sulfophenyl)-2H-tetrazolium, inner salt; MTS(a)] that is bioreduced by cells to form a coloured formazan product. Formation of the formazan product can be measured by absorbance at 490 nm. A media-only “blank” control was included for each time point, which was subtracted from sample absorbance values of the same time point. Absorbance of dTAG-13-treated samples was then normalised to DMSO treatment. MG132 treatment (40 μM) was included as a positive control for each cell line and each time point. Experiment included three technical replicates for each condition, with three separate biological replicates also being conducted. Data was analysed using Excel (Microsoft) and GraphPad Prism software (Version 8).

### Mass spectrometry global proteome and phospho-proteome analysis

Cells were lysed in urea lysis buffer (8 M urea, 50 mM Triethylammonium bicarbonate buffer (TEAB) pH 8.0, supplemented with 1 tablet of cOmplete protease inhibitors per 25 mL lysis buffer and 1 tablet of PhosSTOP phosphatase inhibitors per 10 mL lysis buffer) by Bioruptor® sonication for 15 cycles at 30 sec intervals in LoBind Eppendorf tubes. Lysates were clarified by centrifugation for 20 min at 13,000 G at 4°C. Protein concentration was estimated using the Pierce™ bicinchoninic acid (BCA) method. Equal protein from each condition was reduced with 5 mM dithiothreitol (DTT) at room temperature for 30 min and alkylated with 20 mM iodoacetamide (IAA) in the dark at room temperature for 15 min. Samples were then digested with Lys-C (1:100) at 30°C for 4 hr. Samples were then diluted with 50 mM Triethylammonium bicarbonate buffer (TEAB) to a urea concentration of 1.5 M and were then digested with trypsin (1:20) at 30°C for 16 hr. The digestion was quenched with the addition of trifluoroacetic acid (TFA) to give a final concentration of 1% TFA (v/v) and samples were desalted on 200 mg SepPak C18 cartridges (Waters). For SepPak clean-up, the following solvents were prepared fresh: activation solvent (100% (v/v) acetonitrile (AcN)); Solvent-1 (0.1% (v/v) TFA); Solvent-2 (0.1& (v/v) formic acid (FA)); Solvent-3 (50% (v/v) AcN in 0.1% (v/v) FA). SepPak cartridges were equilibrated with 5 mL 100% AcN, followed by 5 mL 50% MeCN, 0.1% FA and finally with 5 mL 0.1% TFA twice. Samples were then loaded onto the equilibrated C18 cartridges, washed with 5 mL 0.1% TFA four times, followed by washing with 5 mL 0.1% FA. Samples were then eluted with 6 mL 50% AcN, 0.1% FA. Desalted samples were then dried to completeness in a SpeedVac concentrator.

Peptides were resuspended in 50 mM TEAB and labelled using tandem mass tag (TMT) labels as per the manufacturer’s instructions. TMT labels were resuspended in anhydrous acetonitrile, added to assigned samples and incubated for 1 hr at room temperature. Peptides derived from DMSO-treated controls were labelled with TMT labels 126, 127N and 127C, while peptides derived from dTAG-13-treated cells were labelled with 128N, 128C and 129N. Following label check by LC-MS/MS, the labelling reaction was quenched with 5% hydroxylamine for 15 min at room temperature. Labelled peptides from each condition were pooled together and dried.

Pooled peptides were separated by basic reversed phase (bRP) chromatography fractionation on a C18, 250 x 4.6 mm column, 5 μm, XBridge (Waters, Milford, MA) with flow rate at 500 μL/min with two buffers: buffer A (10 mM ammonium formate, pH 10) and buffer B (80% AcN, 10 mM ammonium formate, pH 10). Peptides were resuspended in 100 μL of buffer A (10 mM ammonium formate, pH10) and resolved on a C18 reverse phase column by applying a non-linear gradient of 7-40%. A total of 96 fractions were collected and concentrated into 24 fractions. 90% was used for immobilized metal affinity chromatography (IMAC)-based phospho-peptide enrichment and the remaining 10% for proteomic analysis. Each concentrated fraction was then dried by SpeedVac.

IMAC beads were prepared from Ni-NTA (nitrilotriacetic acid) superflow agarose beads that were stripped of nickel with 100 mM EDTA and incubated in an aqueous solution of 10 mM iron (III) chloride (FeCl3). Dried peptide fractions were reconstituted to a concentration of 0.5 μg/μL in 80% ACN/0.1% TFA. Peptide mixtures were enriched for phosphorylated peptides with 10 μL IMAC beads for 30 min with end-to-end rotation. Enriched IMAC beads were loaded on Empore C18 silica packed stage tips. Stage tips were equilibrated with methanol followed by 50% AcN/0.1% FA then 1% FA. The beads with enriched peptide were loaded onto C18 stage tips and washed with 80% AcN/0.1% TFA. Phosphorylated peptides were eluted from IMAC beads with 500 mM dibasic sodium phosphate, pH 7.0. Enriched phospho-peptides and peptides were analysed on an Orbitrap Fusion Tribrid mass spectrometer interfaced with Dionex Ultimate 3000 nanoflow liquid chromatography system.

For phospho-peptide analysis, peptides were separated on an analytical column (75 μm x 50 cm, RSLC C18) at a flow rate of 300 nL/min using a step gradient of 5-7% solvent B (90% AcN/0.1% FA) for the first 10 min, followed by 7-30% up to 105 min. The total run time was set to 140 min. The mass spectrometer was operated in a data-dependent acquisition mode. A survey full scan MS (from m/z 375-1500) was acquired in the Orbitrap at a resolution of 120,000 at 200 m/z. The automatic gain control (AGC) target for MS1 was set as 2 x 10^5^ and ion filling time set at 50 ms. The most intense ions with charge state ≥2 were isolated and fragmented using higher collision dissociation (HCD) fragmentation, with 38% normalised collision energy, and detected at a mass resolution of 60,000 at 200 m/z. The isolation window was set at 0.7. The AGC target for MS2 was set as 5 x 10^4^ and ion filling time set at 118 ms, while dynamic exclusion was set for 60 s.

For proteomic analysis, peptides were separated on an analytical column (75 μm x 50 cm, RSLC C18) at a flow rate of 300 nL/min, using a step gradient of 5-7% solvent B (90% AcN/0.1% FA) for the first 10 min, followed by 7-25% up to 70 min and 25-35% up to 70-85 min. The total run time was set to 100 min. The mass spectrometer was operated in a data-dependent acquisition mode in SPS MS3 (FT-IT-HCD-FT-HCD) method. A survey full scan MS (from m/z 400-1400) was acquired in the Orbitrap at a resolution of 120,000 at 200 m/z. The AGC target for MS1 was set as 4 x 10^5^ and ion filling time as 50 ms. The precursor ions for MS2 were isolated using a Quadrupole mass filter at a 0.7 Da isolation width, fragmented using a normalized 32% HCD of ion routing multipole and analysed using ion trap. The top 10 MS2 fragment ions in a subsequent scan were isolated and fragmented using HCD at a 65% normalized collision energy and analysed using an Orbitrap mass analyser at a 50 000 resolution, in the scan range of 100–500 m/z.

The proteomics raw data were searched using Sequest HT search engines with Proteome Discoverer 2.1 (Thermo Fisher Scientific). The following parameters were used for searches: Precursor mass tolerance 10 ppm, Fragment mass tolerance 0.1, Enzyme: trypsin, Mis-cleavage: −2, Fixed modification: carbamidomethylation of cysteine residues and TMT of lysine and N-terminal, Dynamic modification: oxidation of methionine. The data were filtered for 1% PSM, peptide and protein level FDR. Only unique peptides were selected for the quantification.

Phosphopeptide-enriched fractions from each replicate were searched against the Uniprot protein database using the Sequest HT search engines with Proteome Discoverer 2.1 (Thermo Fisher Scientific). A 10 plex TMT reporter ion workflow was loaded and the following search parameters were used: trypsin protease was selected; two missed cleavages were allowed; deamidation of Asn and Gln; oxidation of Met and phosphorylation of Ser/Thr were set as variable modifications; and carbamidomethylation of Cys was set as a fixed modification. The default mass error tolerance for MS1 and MS2 (4 ppm and 20 ppm) was used. A minimum of two unique + razor peptides were selected for the quantification. The data were filtered for 1% PSM, peptide, and protein level FDR. For identification of the phospho-site probability, the ptmRS node was used.

### Quantification and statistical analysis of Western blot data

Western blot densitometry was measured using Image Lab and adjusted relative densities were calculated using Excel (Microsoft). All statistical analyses were performed, and graphs were generated using GraphPad Prism software (Version 8). Statistical details including the exact value of n and any statistical tests performed are stated in the figure legends. Graphs display the mean ± standard deviation (SD) of 3 biological replicates, unless stated otherwise in figure legend.

### Quantification and statistical analysis of proteomic data

The Sequest output protein group text files were processed using Perseus software suite (Tyanova et al., 2016). The data was filtered for any proteins identified only by site, common contaminants and reverse hits and proteins identified with single unique peptides. The reporter ion intensities were log2 transformed and the data was normalized by subtracting the median for each sample independently. Student-T test was performed and permutation-based false discovery rate of 5% was applied to identify the differentially enriched and significant protein groups. For cluster analysis, the multiple T test ANOVA was carried out with Benjamin Hochberg correction FDR 5%.

### Bioinformatics analysis

For motif analysis, 16-mer peptides containing the phosphorylated residue at the centre were extracted from the MaxQuant output file and used for motif analysis using WebLogo (available at: https://weblogo.berkeley.edu/) (Schneider and Stephens, 1990, Crooks et al., 2004). For the comparison of our data with known PPP2CA substrates, we searched our identified substrates against PPP2CA substrate listed on the DEPOD database - the human DEPhOsphorylation Database (available at: http://depod.bioss.uni-freiburg.de/) (Damle and Kohn, 2019). The KinMap beta tool (available at: http://www.kinhub.org/kinmap/index.html) was used to build the kinome map. We plotted the list of kinases identified in our dataset as potential PPP2CA substrates and highlighted them on the kinome map. To identify the upstream kinases whose substrates are overrepresented amongst our identified putative PPP2CA substrates, Kinase Enrichment Analysis 3 (KEA3) was carried out (available at: https://maayanlab.cloud/kea3/#results) (Kuleshov et al., 2021). We used Funrich (Version 3.1.3) (https://www.FunRich.org) (Pathan et al., 2015) and EnrichR (available at: https://maayanlab.cloud/Enrichr) (Kuleshov et al., 2016, Chen et al., 2013) to conduct the KEGG pathway analysis, uncover associations with biological processes, molecular function, localisation, protein domain architecture and diseases. Further protein–protein interaction network analysis for the putative PPP2CA substrates was carried out using STRING (available at: https://string-db.org/) (Snel et al., 2000). The proteins that did not show any connection with the network were removed for clarity. To identify proteins containing the PP2A-specific SLiM, the LSPIxE motif was scanned across the human proteome using FIMO (Find Individual Motif Occurrences) (available at: https://meme-suite.org/meme/tools/fimo) (Grant et al., 2011).

## Resource Availability

All mass spectrometry data acquired from this study has been deposited in the PRIDE database. Additionally, the data can be accessed via the following CurtainPTM (Phung et al., 2023) links: For phospho-proteomics: https://curtainptm.proteo.info/#/7bbd6bee-d15f-415a-8ffe-da9f8bb91873 For total proteomics: https://curtain.proteo.info/#/c16e75ef-879c-4e9a-9186-4bc4b5f28f0a Raw data will be uploaded to Mendeley Data.

## Lead Contact

Further information and requests for resources and reagents should be directed to and will be fulfilled by the Lead Contact, Gopal Sapkota.

## Author Contributions

AB and GS performed experiments, collected and analysed data and contributed to the writing of the manuscript. BEP performed some optimisation experiments. TJM designed the strategies for and generated the CRISPR/Cas9 constructs used in this study. G.P.S. conceived the project, analysed data, and contributed to the writing of the manuscript.

## ACKNOWLEDGEMENTS

GPS is supported by the UKRI Medical Research Council (grant MC_UU_00018/6) and the pharmaceutical companies supporting the Division of Signal Transduction Therapy (Boehringer Ingelheim, GlaxoSmithKline, Merck-Serono). For this study, AB was supported by UKRI BBSRC EASTBIO PhD studentship (grant BB/M010996/1). We thank the Sapkota lab members for critical appraisal of the data. We thank E. Allen, A. Muir, S. Dalglish, E. Webster and J. Stark for help and assistance with tissue culture, the staff at the DNA Sequencing services (School of Life Sciences, University of Dundee), and the cloning and antibody teams within the MRC-PPU Reagents and Services (University of Dundee, coordinated by J. Hastie. We thank the staff at the flow cytometry facility (School of Life Sciences, University of Dundee) for their invaluable help and advice throughout this project.

## Supplementary Figure Legends

**Figure S1.**
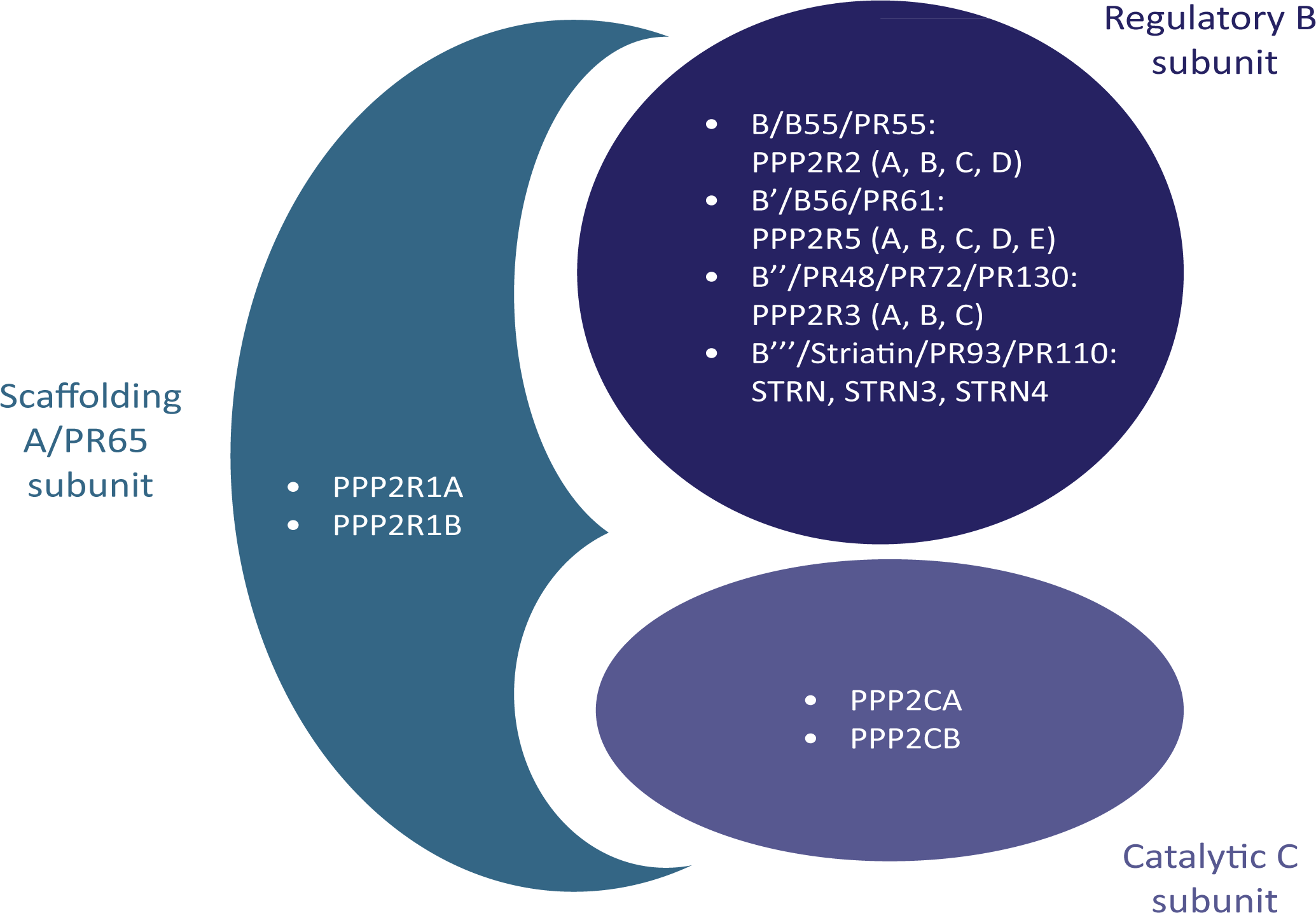
Human PP2A phosphatase holoenzyme structure with genes encoding different subunit isoforms. PP2A holoenzyme complex detailing genes encoding distinct isoforms of the scaffolding A, regulatory B and catalytic C subunits.

**Figure S2.**
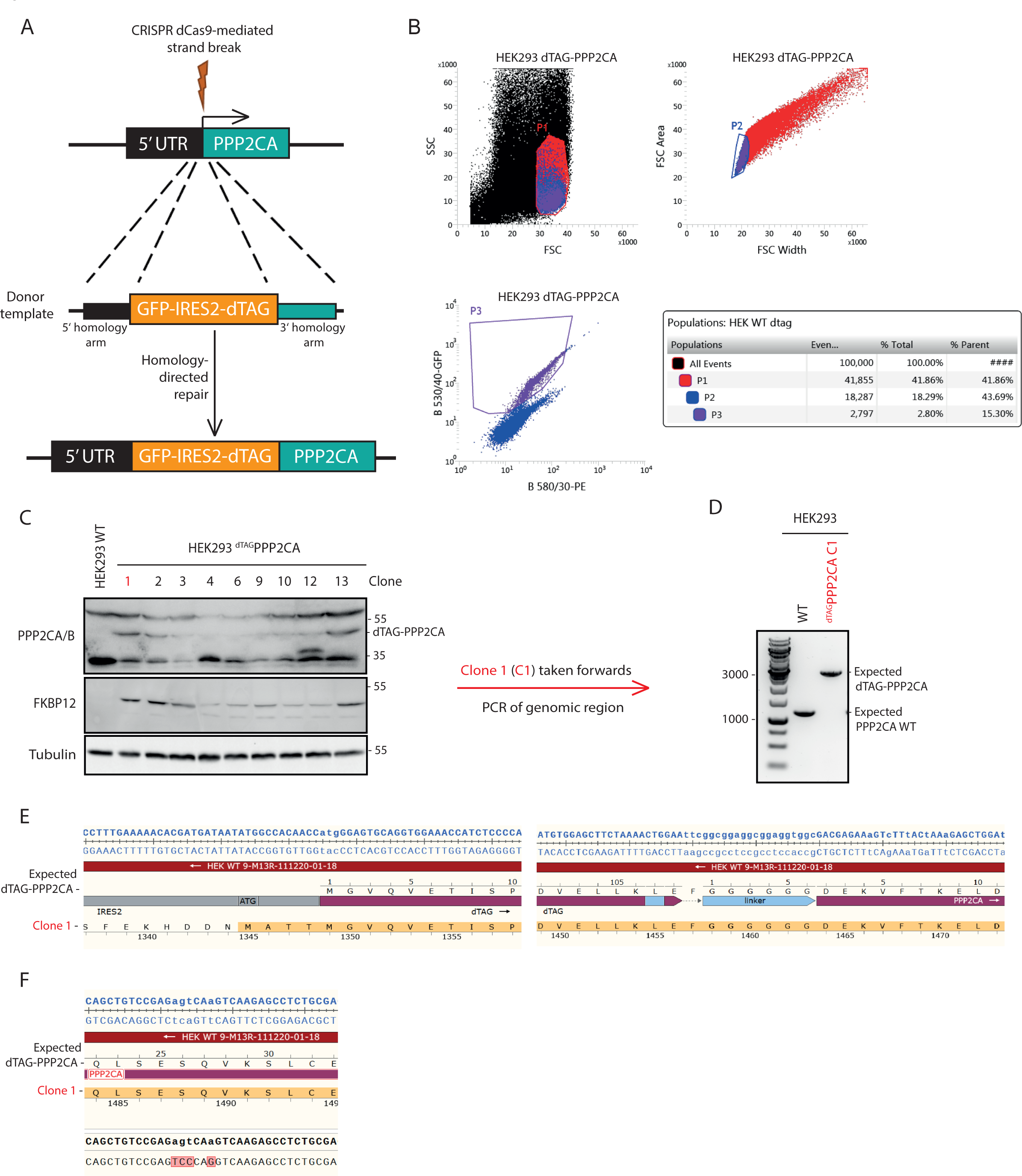
Confirmation of ^dTAG/dTAG^PPP2CA knock-in in HEK293 cells. **A.** Depiction of CRISPR strategy to introduce dTAG at N-terminus of PPP2CA. **B.** Fluorescence-activated cell sorting (FACS) analysis of HEK293 GFP-IRES2-dTAG-PPP2CA cells identified 15.30% of cells as being GFP-positive. **C.** Initial immunoblot screen of isolated single cell clones from 24-well plates identified clone 1, among others, as potential homozygous for dTAG-PPP2CA knock-in. Anti-FKBP12 antibody was used to detect dTAG (FKBP12^F36V^). **D.** Confirmation of knock-in by polymerase chain reaction (PCR), using PPP2CA forward (Fw) and reverse (Rev) primers that bind upstream of the 5’-UTR region and downstream of the start of the PPP2CA gene to give a PCR product of ∼2.8 kbp for dTAG-PPP2CA or a product of ∼1.2 kbp for wild type (WT) PPP2CA. Clone 1 was identified as containing homozygous knock-in of ^dTAG/dTAG^PPP2CA. **E.** Genomic DNA sequencing and subsequent alignment with the expected DNA sequence confirmed successful knock-in at IRES2-dTAG and dTAG-PPP2CA boundaries, respectively. Alignment conducted using SnapGene. **F.** Sequence alignment of genomic DNA extract from ^dTAG/dTAG^PPP2CA HEK293 Clone 1 cells identified expected silent mutations at residues 26 and 27 that were introduced on the donor to block recognition by the guide RNAs. Alignment conducted using SnapGene.

**Figure S3.**
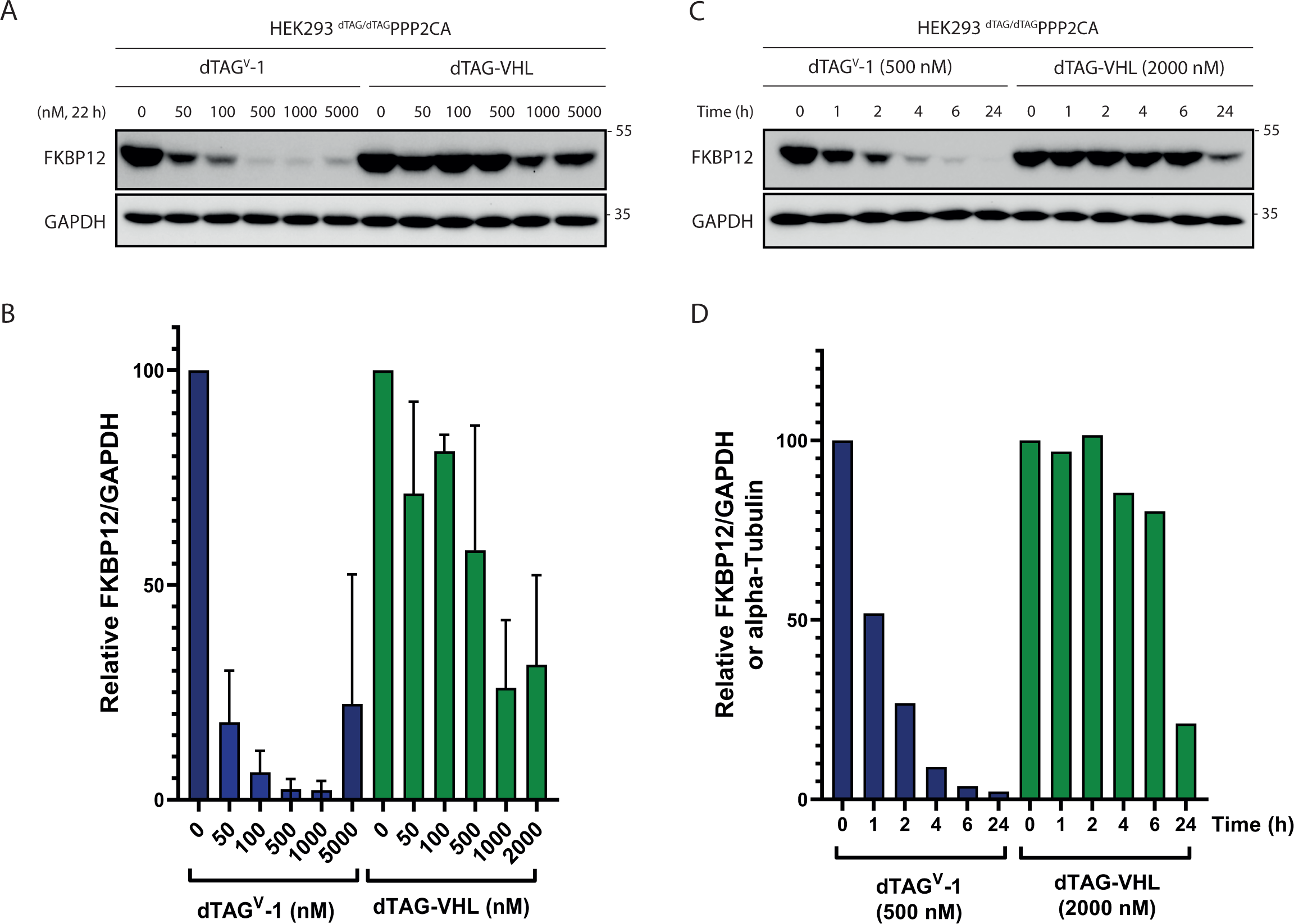
Degradation of dTAG-PPP2CA using dTAG-^V^1 and dTAG-VHL PROTACs. **A-B.** ^dTAG/dTAG^PPP2CA HEK293 cells were treated with the indicated concentrations of dTAG^V^-1, dTAG-VHL or DMSO for 22 h prior to lysis and immunoblot analysis. Anti-FKBP12 antibody was used here to detect dTAG (also known as FKBP12^F36V^). Three biological replicates are quantified in **B**, where mean values are displayed for normalised (FKBP12/GAPDH), relative to DMSO-treated control samples. Error bars represent standard deviation. **C-D.** ^dTAG/dTAG^PPP2CA HEK293 cells were treated with 500 nM dTAG^V^-1 or 2000 nM dTAG-VHL for indicated durations prior to lysis and immunoblot analysis. Quantification of two biological replicates in **D**, with mean values shown, relative to 0 h treatment.

**Figure S4.**
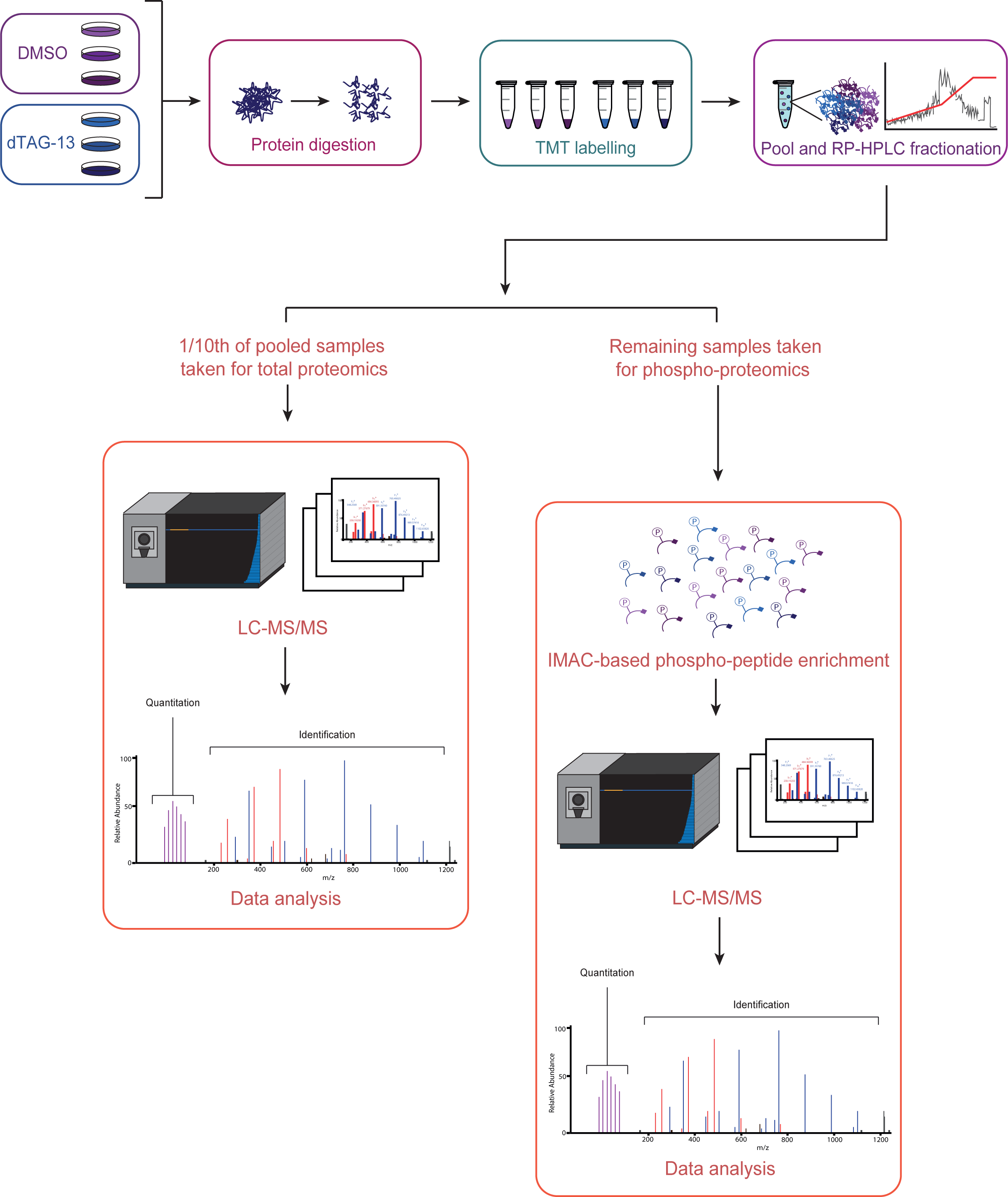
Global total- and phospho-proteomic approach to explore substrate landscape of PPP2CA. Workflow of total- and phospho-proteomic sample preparation and analysis using ^dTAG/dTAG^PPP2CA HEK293 cells treated with dTAG-13 (100 nM) or DMSO for 24 h.

**Figure S5.**
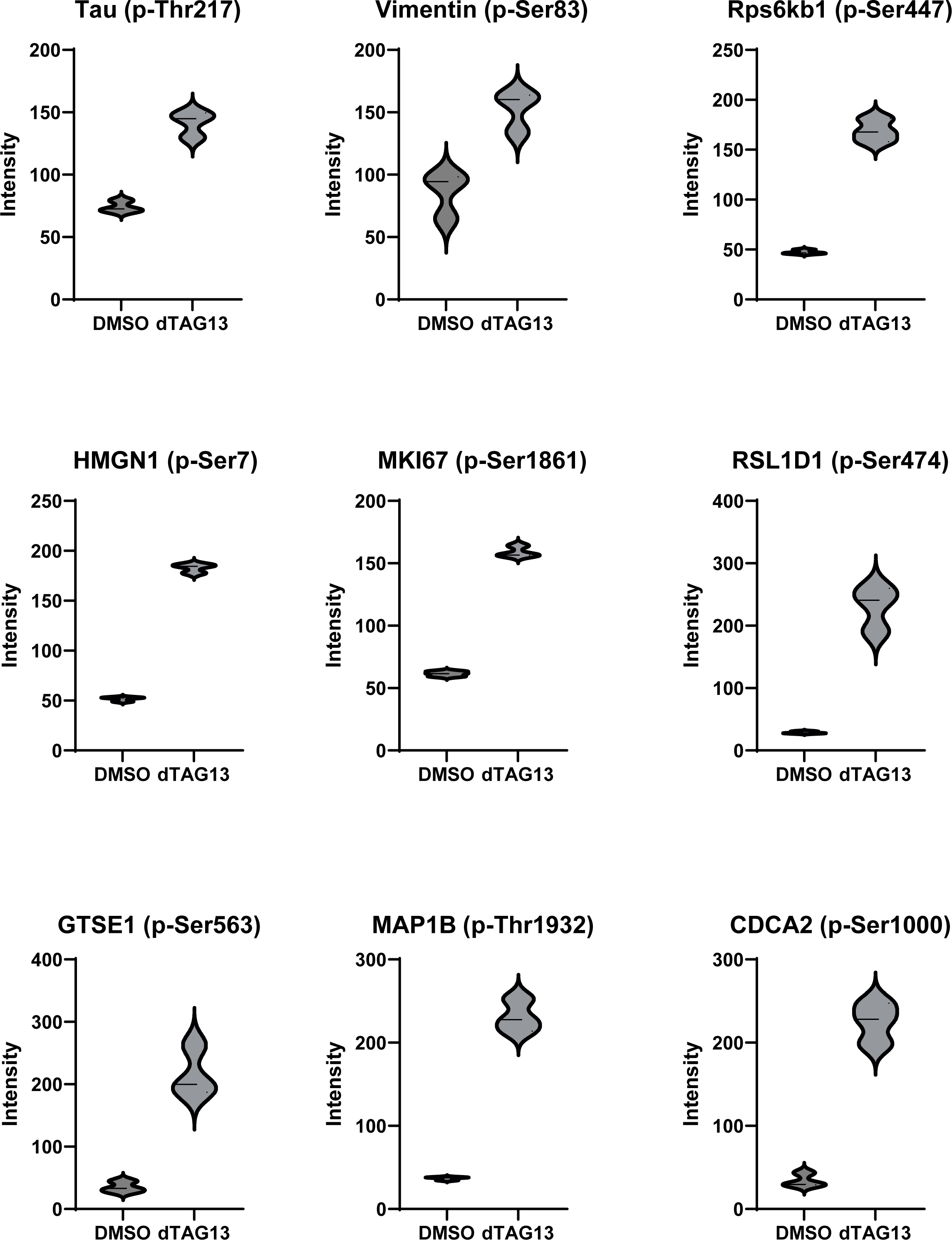
Violin plots of selective phospho-peptides identified from quantitative phospho-proteomic analysis. Some phospho-peptides shown here have been validated by immunoblotting in Figure 5.

**Figure S6.**
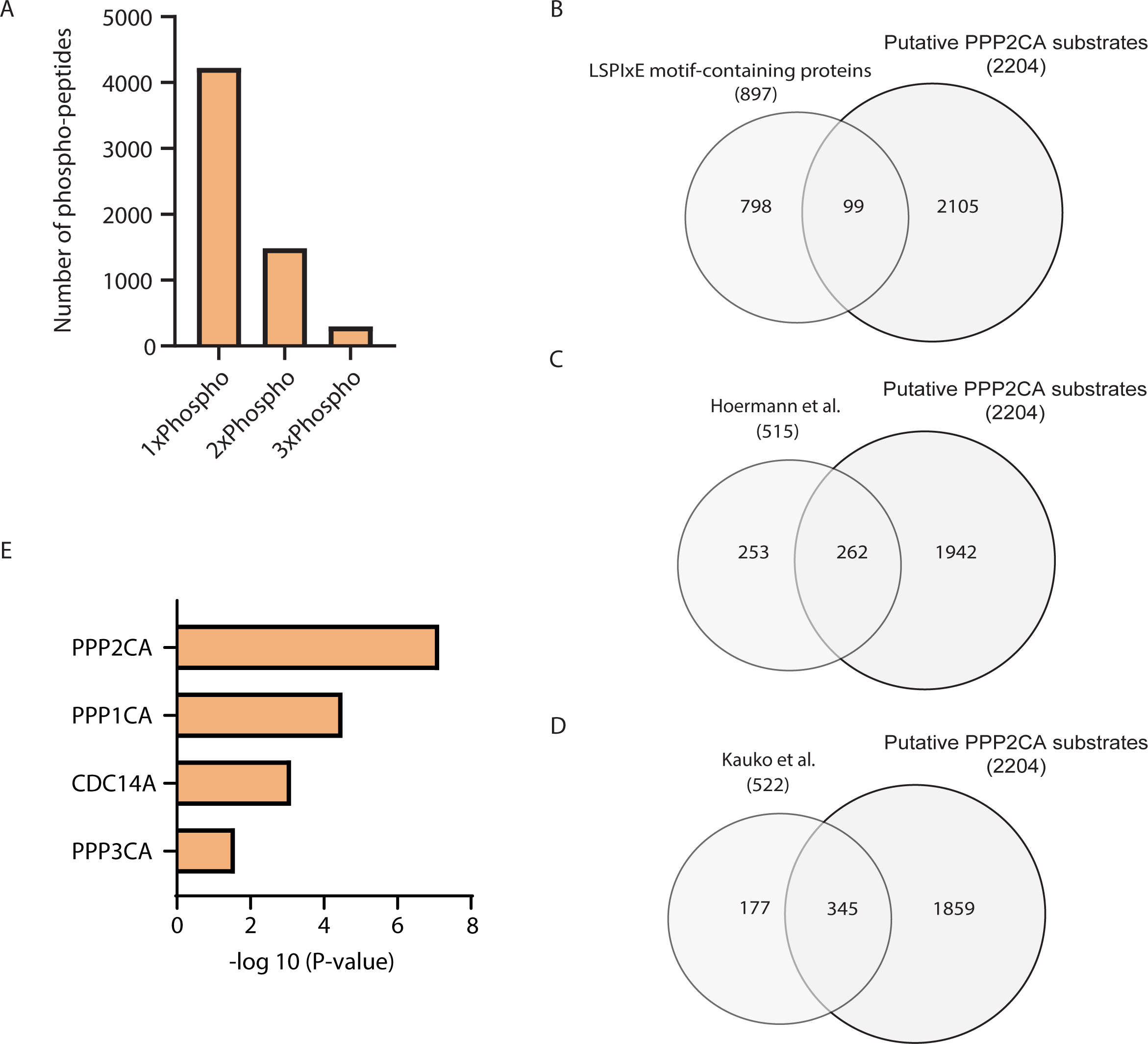
Analysis of putative PPP2CA substrates. **A.** The prevalence of mono-, di- and tri-phosphorylated peptides amongst the phospho-peptides we identified to be enriched upon dTAG-PPP2CA degradation. **B.** PP2A-B56α-specific SLiM LSPIxE was scanned across the human proteome using Find Individual Motif Occurrences (FIMO) (available at: https://meme-suite.org/meme/tools/fimo). This identified 897 unique proteins containing the LSPIxE SLiM. We compared these with our identified putative PPP2CA substrates, with common proteins displayed in the Venn diagram. **C.** Comparison between PPP2CA substrates identified by Hoermann et al. (2020) and putative PPP2CA substrates identified in our study. **D.** Comparison between PPP2CA substrates identified by Kauko et al. (2020) and putative PPP2CA substrates identified in our study. **E.** Using Enrichr, putative PPP2CA substrates identified in our study were compared with the DEPOD database to uncover phosphatases the database predicted to be most likely to dephosphorylate these substrates (available at: https://maayanlab.cloud/Enrichr).

**Figure S7.**
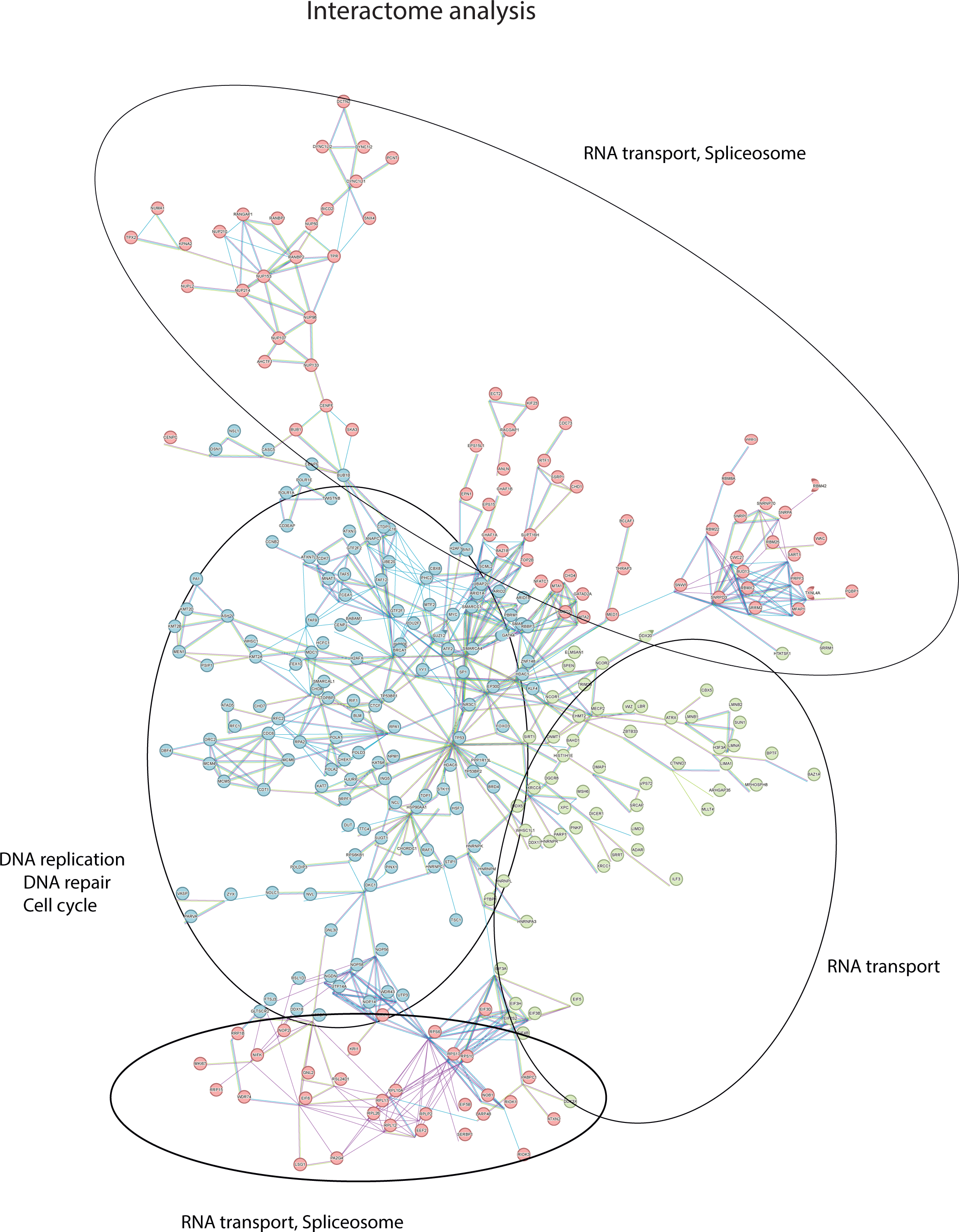
Network analysis of phospho-proteins identified as putative PPP2CA substrates from the phospho-proteomic analysis. Four major enriched pathways are circled and presented. The colour of the nodes indicates the biological process that the proteins in the network are involved in.

## SUPPLEMENTARY TABLES

**Table S1:** Full list of total proteomics data.

**Table S2:** Full list of phospho-proteomics data, including separate sheets for Significant, Hyperphosphorylated and Hypophosphorylated hits.

**Table S3:** Full lists of results from different bioinformatic analyses conducted in this study using the hyperphosphorylated hits are detailed in separate sheets, including for KEGG pathway analysis, biological processes, molecular functions, protein domains, subcellular localisation, disease associations and comparisons with DEPOD database.

